# Two type II phosphatidylinositol 4-kinases function sequentially in tubule-mediated cargo delivery from early endosomes to melanosomes

**DOI:** 10.1101/2021.10.19.465056

**Authors:** Yueyao Zhu, Shuixing Li, Alexa Jaume, Riddhi Atul Jani, Cédric Delevoye, Graça Raposo, Michael S. Marks

## Abstract

Melanosomes are pigment cell-specific lysosome-related organelles in which melanin pigments are synthesized and stored. Melanosome maturation requires delivery of melanogenic cargoes via tubular transport carriers that emanate from early endosomes and that require BLOC-1 for their formation. Here we show that phosphatidylinositol-4-phosphate (PtdIns4P) and the type II PtdIns-4-kinases (PI4KIIα and PI4KIIβ) support BLOC-1-dependent tubule formation to regulate melanosome biogenesis. Depletion of either PI4KIIα or PI4KIIβ with shRNAs in melanocytes reduced melanin content and misrouted BLOC-1-dependent cargoes to late endosomes/lysosomes. Genetic epistasis, cell fractionation, and quantitative live-cell imaging analyses show that PI4KIIα and PI4KIIβ function sequentially and non-redundantly downstream of BLOC-1 during tubule elongation towards melanosomes by generating local pools of PtdIns4P. The data show that both type II PtdIns-4-kinases are necessary for efficient BLOC-1-dependent tubule elongation and subsequent melanosome contact and content delivery during melanosome biogenesis. The independent functions of PtdIns-4-kinases in tubule extension are downstream of likely redundant functions in BLOC-1-dependent tubule initiation.

**SUMMARY:** Contents are delivered to maturing melanosomes from early endosomal intermediates through tubular transport carriers. Zhu et al show that two type II phosphatidylinositol kinases, PI4KIIα and PI4KIIβ, sequentially generate phosphatidylinositol-4-phosphate during tubule initiation and elongation for ultimate melanosome content delivery.

## INTRODUCTION

The endolysosomal system is a dynamic network of intracellular membrane-bound compartments that plays an essential global role in nutrient uptake, signaling, metabolism, and pathogen life cycle (Huotari and Helenius, 2011; Klumperman and Raposo, 2014). In some cell types, the endolysosomal system is additionally adapted to create specialized subcellular compartments termed lysosome-related organelles (LROs) for specific physiological functions (Bowman et al., 2019; Delevoye et al., 2019). LROs share some features with conventional lysosomes but possess distinct morphological and functional characteristics that are conferred largely by cell type-specific contents (Bowman et al., 2019; Delevoye et al., 2019). Many LROs share common origins, and abnormalities in LRO biogenesis in several cell types underlie the pathology of the Hermansky–Pudlak syndromes (HPS), a group of inherited multisystem disorders characterized by oculocutaneous albinism, bleeding diathesis, and other variable symptoms. HPS symptoms reflect defective biogenesis of melanosomes in pigment cells, dense granules in platelets, and additional LROs in other cell types (Bowman et al., 2019; De Jesus Rojas and Young, 2020). How the products of HPS-associated genes and their interactors orchestrate LRO biogenesis is only beginning to be understood.

The genes that are defective in HPS encode essential subunits of four protein complexes – adaptor protein-3 (AP-3) and <u>b</u>iogenesis of lysosome-related <u>o</u>rganelles <u>c</u>omplex (BLOC)-1, −2 and −3 – that regulate membrane dynamics essential for the biosynthetic delivery and/or recycling of components to or from maturing LROs (Bowman et al., 2019; Di Pietro and Dell’Angelica, 2005). BLOC and AP-3 functions in LRO biogenesis are best understood for melanosome maturation in epidermal melanocytes (Le et al., 2021). Melanosomes mature from nonpigmented stage I and II precursors to pigmented stage III and IV organelles through the delivery of melanogenic enzymes and transporters (Novikoff et al., 1968; Raposo et al., 2001; Seiji et al., 1961) via three district pathways: one from the Golgi (Patwardhan et al., 2017), and two from early endosomes that are dysregulated in HPS (Bowman et al., 2019). One of these pathways uniquely requires BLOC-1 to generate recycling-endosome-like tubular transport intermediates from early sorting endosomes that ultimately fuse with melanosomes, delivering cargoes such as tyrosinase-related protein-1 (TYRP1) and the melanosomal chloride channel (Bellono and Oancea, 2014; Sitaram et al., 2009) oculocutaneous albinism type 2 (OCA2) (Delevoye et al., 2009; Setty et al., 2008; Setty et al., 2007; Sitaram et al., 2012). BLOC-1 coordinates with the microtubule motor KIF13A and actin polymerization to generate the tubules (Delevoye et al., 2016; Delevoye et al., 2009) and may directly stabilize them (Jani et al., 2021; Lee et al., 2012). Adaptor protein-1 (AP-1) and a BLOC-1-associated cohort of AP-3 sorts cargoes and SNARE proteins (including the v-SNARE, VAMP7) into the melanosome-bound tubules (Bowman et al., 2021; Delevoye et al., 2009; Dennis et al., 2015; Theos et al., 2005). AP-1 also binds to and might recruit KIF13A to the tubules (Delevoye et al., 2009; Delevoye et al., 2014; Nakagawa et al., 2000). Despite these advances, the field lacks a comprehensive mechanistic understanding of BLOC-1-dependent tubule formation. In particular, roles for lipids in the process are unknown.

Among lipids, phosphoinositides are good candidates to regulate BLOC-1-dependent tubule formation. Phosphoinositides function in determining membrane identity, spatio-temporal regulation of membrane trafficking, and membrane remodeling, often by binding proteins that induce or stabilize membrane deformation (Balla, 2013; Suetsugu et al., 2014). In particular, phosphatidylinositol 4-phosphate (PtdIns4P) controls membrane curvature associated with membrane tubules in the endolysosomal system (Levin-Konigsberg et al., 2019; López-Haber et al., 2020; Ma et al., 2020; McGrath et al., 2021; Rahajeng et al., 2019). The cellular distribution of PtdIns4P is governed by the balanced activities of PtdIns4P-generating kinases and phosphoinositide phosphatases (D’Angelo et al., 2008; Tan and Brill, 2014). The endosomal PtdIns4P pool is largely produced by the type II PtdIns-4-kinases, PI4KIIα and PI4KIIβ (Balla and Balla, 2006; Minogue, 2018). Type II PtdIns-4-kinases and PtdIns4P are required for Weibel-Palade body maturation in endothelial cells (Lopes da Silva et al., 2016) and to form tubules during phagolysosome resolution in macrophages (Levin-Konigsberg et al., 2019), phagosome signaling in dendritic cells (López-Haber et al., 2020), and secretory granule maturation in *Drosophila melanogaster* larval salivary glands (Burgess et al., 2012; Ma et al., 2020). These findings suggest a general role for type II PtdIns-4-kinases and PtdIns4P in LRO biogenesis.

Their association with membrane tubules suggests that PtdIns4P and PtdIns-4-kinases might function in BLOC-1-dependent transport during LRO biogenesis. Accordingly, PI4KIIα binds directly to AP-3 (Craige et al., 2008; Salazar et al., 2005) and indirectly to BLOC-1 (Salazar et al., 2009), and requires AP-3 for targeting to lysosomes and LROs in other cell types (Craige et al., 2008; López-Haber et al., 2020). Similarly, PI4KIIβ binds directly to AP-1 (Wieffer et al., 2013). Moreover, BLOC-1 binds to PtdIns4P and other phosphoinositides on tubular membranes in vitro, and requires type II PtdIns-4-kinases to initiate recycling endosome formation in HeLa cells (Jani et al., 2021). Here, we used gene silencing together with imaging and biochemical approaches to test whether PI4KIIα and/or PI4KIIβ function in BLOC-1-dependent cargo transport to melanosomes by generating PtdIns4P on endosomal tubules. While PI4KIIα and PI4KIIβ likely function redundantly in BLOC-1-dependent tubule initiation (Jani et al., 2021), we show that they play additional non-redundant roles in generating PtdIns4P on the tubular carriers in melanocytes. This is necessary for tubule stability and for cargo delivery to melanosomes.

## RESULTS

### Type II PtdIns4-kinases are required for melanosomal localization of BLOC-1-dependent cargos

The biosynthetic delivery of TYRP1 to melanosomes from early endosomes in melanocytes requires BLOC-1 (Setty et al., 2007). Thus, to determine if type II PtdIns-4-kinases function in BLOC-1-dependent cargo transport, we first tested whether depletion of PI4KIIα or PI4KIIβ impacts TYRP1 localization. Darkly pigmented immortalized wild-type (WT) melan-Ink4a melanocytes (Sviderskaya et al., 2002) were transduced with lentiviruses to co-express puromycin resistance and shRNA for PI4KIIα (shPI4KIIα), PI4KIIβ (shPI4KIIβ), the core Pallidin subunit of BLOC-1 (shPallidin), or a non-target control (shNC), and puromycin-resistant cells were selected for 7-9 d. Two distinct shRNAs per target each effectively reduced target expression but not other proteins analyzed (**Fig. S1A-C**). TYRP1 steady state localization was assessed by confocal immunofluorescence microscopy (IFM) relative to pigment granules visualized by bright field microscopy (**Fig. 1A-C**); quantitative colocalization analyses excluded the crowded nuclear and perinuclear regions in which individual compartments could not be distinguished. As in untreated cells (Raposo et al., 2001; Setty et al., 2007; Vijayasaradhi et al., 1995), the majority of peripheral TYRP1 in shNC cells (**Fig. 1Aa-c, 1B, 1C**) surrounded pigment granules, apparent by IFM as fluorescent “donuts” filled with black melanin (white arrows) and representing localization to the membrane of mature melanosomes (Raposo et al., 2001). In contrast, TYRP1 in shPI4KIIα (**Fig. 1Ad-f**) or shPI4KIIβ cells (**Fig. 1Ag-i**) localized to punctate structures that largely did not overlap with pigment granules (yellow arrowheads), reducing peripheral melanosome localization by ∼50% (**Fig. 1B**). Moreover, many shPI4KIIβ and particularly shPI4KIIα cells were hypopigmented (**Fig. 1Ab, Ae, Ah**), most apparent by 14 d after transduction (**Fig. S2A**) and confirmed by a quantitative melanin content assay (**Fig. 1F**; the low pigmentation induced by shPI4KIIα-2 may reflect an off-target effect because it also slowed cell growth and depleted PI4KIIα less effectively than shPI4KIIα-1). Nevertheless, the fraction of remaining melanosomes labeled by TYRP1 in shPI4KIIα and shPI4KIIβ cells was reduced by ∼50% relative to shNC cells (**Fig. 1C**). These effects were specific to PI4KIIα and PI4KIIβ, as depletion of the type III PtdIns4P kinase, PI4KIIIβ, did not affect TYRP1 localization or pigmentation (**Fig. S1D-G**). Impaired TYRP1 localization is also observed in mouse melanocytes lacking BLOC-1 subunits (Setty et al., 2007); indeed, in highly depigmented BLOC-1-deficient shPallidin cells, TYRP1 overlap with remaining pigment granules was similar to that of shPI4KIIα or shPI4KIIβ cells (**Figs. 1Aj-l, 1B, S1B, S1C**). Thus, like BLOC-1, both PI4KIIα and PI4KIIβ but not PI4KIIIβ are required for TYRP1 localization to melanosomes and optimal pigmentation.

**Figure 1.**
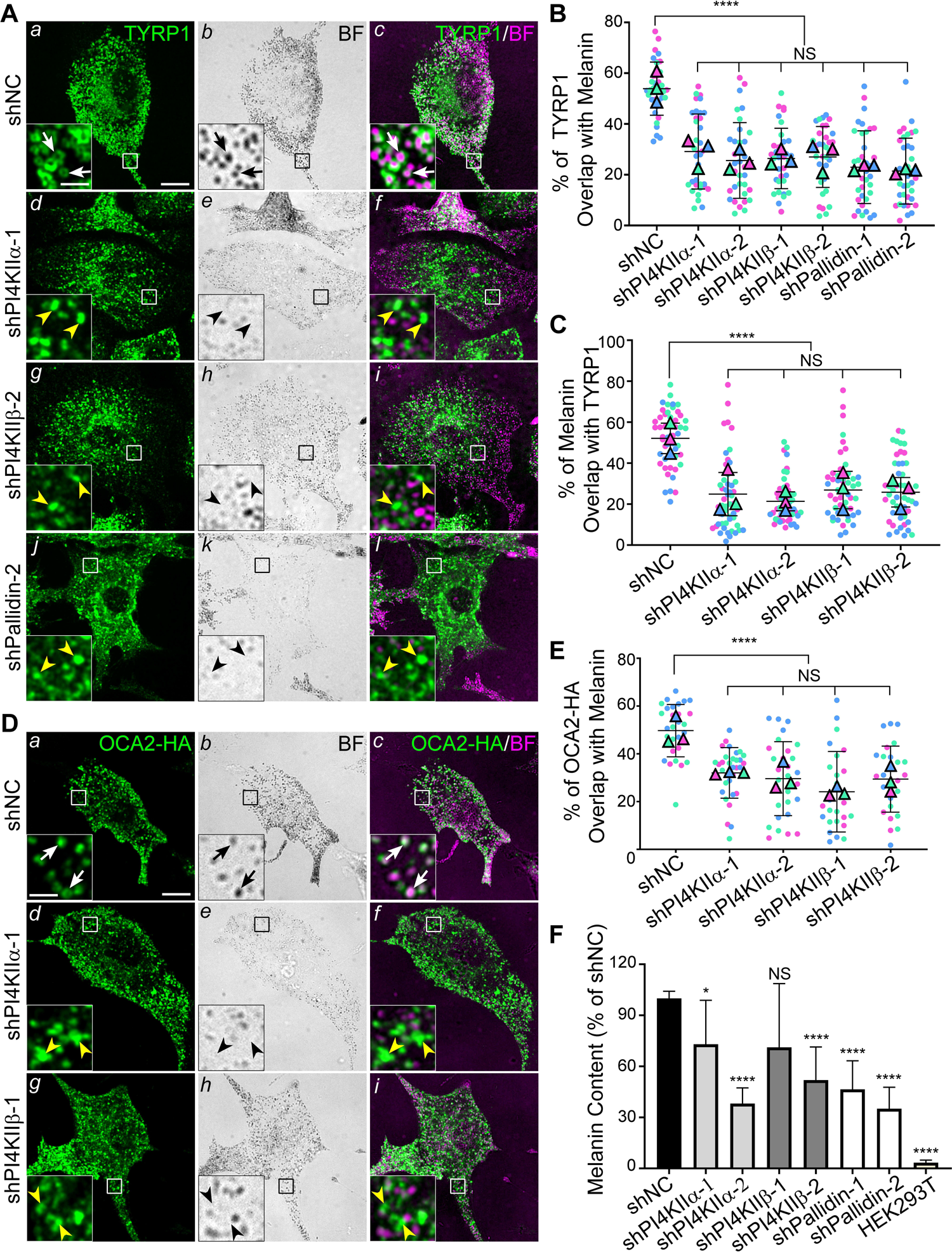
Depletion of either PI4KIIα or PI4KIIβ by shRNA in WT melanocytes impairs localization of BLOC-1-dependent cargoes to melanosomes. WT melan-Ink4a melanocytes were transduced with lentiviruses to express shNC or the indicated shRNAs to PI4KIIα, PI4KIIβ, or the Pallidin subunit of BLOC-1 and selected for 7-9 d (A-E) or 14 d (F). Cells in D, E were additionally transiently transfected to express HA-tagged OCA2 (OCA2-HA) 2 d prior to analysis. **(*A* and *D*)** Cells were analyzed by IFM for TYRP1 (A; green, left and right panels) or HA (D; left and right panels), and by bright field microscopy (BF) for pigment granules (middle; pseudocolored magenta at right). White arrows, pigment granules surrounded by TYRP1 or overlapping with OCA2-HA; yellow arrowheads, TYRP1 or OCA2-HA not associated with pigment granules. Boxed regions are magnified 5-fold in insets. Scale bars: main panels, 10 μm; insets, 2 μm. **(*B, C and E*)** Quantification of TYRP1 (B) or HA-OCA2 (E) overlap with melanosomes or melanosome coating by TYRP1 (C). Data from three independent experiments each were analyzed by Kruskal-Wallis (B and C) or Welch’s ANOVA (E); ****, p<0.0001; NS, not significant. **(*F*)** shRNA-treated melan-Ink4a cells or HEK293T cells as a negative control were analyzed by quantitative melanin content assay. Data are presented as mean ± SD of five independent experiments as a percentage of the signal from shNC-treated cells, and were analyzed by Welch’s ANOVA relative to shNC-treated cells; ****, p<0.0001; *, p<0.05; NS, not significant.

We considered generating PI4KIIα and PI4KIIβ gene knockouts, but in shPI4KIIα and shPI4KIIβ melanocytes at 30 d post-infection, TYRP1 localization to melanosomes was largely restored to control levels despite consistent target protein depletion (**Fig. S2B**, white arrows; **Figs. S2C, D**). This likely reflects functional compensation for the loss of each kinase over time. All subsequent analyses were thus performed on cells expressing shRNAs for 7-9 d.

To test whether PI4KIIα or PI4KIIβ depletion impacts other BLOC-1-dependent melanosome cargoes (Dennis et al., 2016; Sitaram et al., 2012), we analyzed the localization of transiently expressed, epitope-tagged forms of OCA2 and VAMP7 relative to melanosomes in shPI4KIIα and shPI4KIIβ cells. Compared with shNC cells in which HA-tagged OCA2 (OCA2-HA; **Figs. 1Da-c, 1E**) and GPF-tagged VAMP7 (GFP-VAMP7; **Fig. S3Aa-c, S3B**) largely overlapped with pigment granules (white arrows), localization of both cargoes to melanosomes was impaired in shPI4KIIα and shPI4KIIβ cells (**Figs. 1Dd-i, 1E, S3Ad-i, S3B)**. Like TYRP1, mislocalized OCA2-HA and GFP-VAMP7 accumulated in punctate structures throughout the cytoplasm (**Figs. 1D, S3A**, yellow arrowheads). Thus, both type II PtdIns-4-kinases in melanocytes are required for effective BLOC-1-dependent cargo delivery.

### TYRP1 is largely mislocalized to late endosomes and lysosomes in cells depleted of either PI4KIIα or PI4KIIβ

BLOC-1 is required for TYRP1 and other cargoes to exit vacuolar endosomes into tubules destined for maturing melanosomes, and thus these cargoes are trapped in enlarged early endosomes in BLOC-1-deficient melanocytes (Delevoye et al., 2016; Dennis et al., 2016; Setty et al., 2008; Setty et al., 2007; Sitaram et al., 2012). To determine whether PI4KIIα and PI4KIIβ function at the same step as BLOC-1, shNC, shPI4KIIα, shPI4KIIβ, and shPallidin cells were fixed, immunolabeled for TYRP1 and the pan-early endosomal t-SNARE syntaxin-13 (STX13), and analyzed by IFM. Whereas STX13 and TYRP1 overlapped minimally in shNC cells (**Fig. 2Aa-c**, white arrows; **Fig. 2B**), they overlapped extensively in the periphery of shPallidin cells (**Fig. 2Aj-l**, yellow arrowheads; **Fig. 2B**) as expected for cells lacking BLOC-1 subunits (Setty et al., 2007). Although the overlap of TYRP1 with STX13 (yellow arrowheads) in shPI4KIIα and shPI4KIIβ cells was higher than in shNC cells, it was not as high as in shPallidin cells (**Fig. 2Ad-i**; **Fig. 2B**). These results indicate that while depletion of either PI4KIIα or PI4KIIβ partially traps TYRP1 in early endosomes, consistent with a redundant requirement for either kinase to initiate BLOC-1-dependent cargo export from endosomes (Jani et al., 2021), neither alone is absolutely required for BLOC-1-dependent cargo exit from early sorting endosomes.

**Figure 2.**
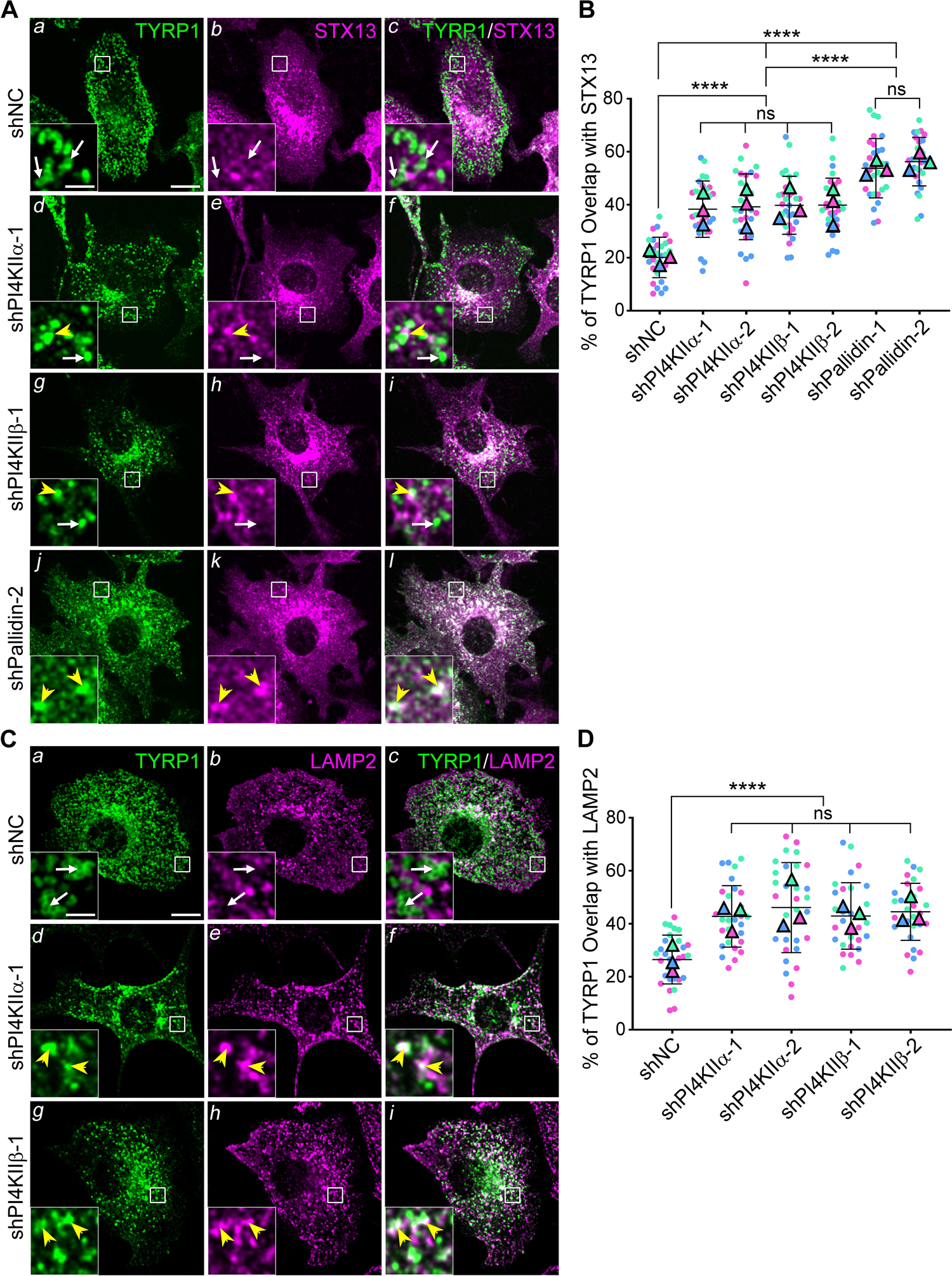
TYRP1 is mislocalized to early endosomes and late endosomes/ lysosomes in WT melanocytes depleted of either PI4KIIα or PI4KIIβ. Melan-Ink4a melanocytes were transduced with lentiviruses to express shNC or the indicated shRNAs to PI4KIIα, PI4KIIβ, or the Pallidin subunit of BLOC-1, and selected for 7 to 9 d. **(*A* and *C*)** Cells were analyzed by IFM for TYRP1 (green, left and right panels) relative to STX13 to mark early endosomes (A) or LAMP2 to mark late endosomes and lysosomes (C) (magenta, middle and right panels). White arrows, TYRP1 not associated with STX13 or LAMP2; yellow arrowheads, TYRP1 overlapping with STX13 or LAMP2. Boxed regions are magnified 5-fold in the insets. Scale bars: main panels, 10 μm; insets, 2 μm. **(*B* and *D*)** Quantification of TYRP1 overlap with STX13 (B) or LAMP2 (D). Data from three independent experiments were analyzed by ordinary one-way ANOVA (B) or Welch’s ANOVA (D); ****, p<0.0001; ns, not significant.

To determine where most TYRP1 accumulated in shPI4KIIα and shPI4KIIβ cells, we conducted additional IFM analyses. Notably, whereas overlap of peripheral TYRP1 with the late endosomal/ lysosomal membrane protein, LAMP2, is minimal in shNC cells (**Fig. 2Ca-c**, white arrows; **Fig. 2D**) – as also observed in untreated WT or BLOC-1-deficient cells (Setty et al., 2007) – LAMP2 overlap was substantially increased (yellow arrowheads) in shPI4KIIα and shPI4KIIβ cells (**Fig. 2Cd-i**; **Fig. 2D**). These data indicate that in the absence of PI4KIIα or PI4KIIβ, a cohort of TYRP1 exits early endosomes but is then mistargeted to late endosomes and lysosomes. Thus, both type II PtdIns4-kinases have non-redundant functions in cargo delivery to melanosomes in a separate and likely subsequent step from BLOC-1.

### PI4KIIα and PI4KIIβ functions in TYRP1 trafficking require their lipid kinase activity and binding to adaptor proteins

To ensure that cargo mislocalization in shPI4KIIα and shPI4KIIβ cells reflected target depletion, we assessed whether TYRP1 localization to melanosomes was restored by overexpressing shRNA-resistant PI4KIIα or PI4KIIβ. WT mouse melan-Ink4a melanocytes were transduced with retroviruses encoding GFP-tagged human WT PI4KIIα or PI4KIIβ, selected for stable integration for at least two weeks, and then depleted of the endogenous target protein by mouse-specific shPI4KIIα or shPI4KIIβ. Cells were analyzed 7-9 d later by IFM and bright field microscopy. Whereas TYRP1 overlapped minimally with pigment granules in melanocytes expressing shPI4KIIα or shPI4KIIβ alone (yellow arrowheads, **Figs. 3B, 4B**), co-expression of PI4KIIα-WT-GFP with shPI4KIIα-1 (**Fig. 3C**) or of PI4KIIβ-WT-GFP with shPI4KIIβ-2 (**Fig. 4C**) restored a WT pattern of TYRP1 around pigment granules in a significant fraction of cells (compare white arrows in **Fig. 3A** with **3C** and **Fig. 4A** with **4C**; **Figs. 3G, 4G**). Immunoblotting showed that the shRNAs efficiently reduced target expression even in the presence of correctly expressed excess human GFP-tagged transgenes (**Figs. 3H, 4H**). Thus, the impaired TYRP1 localization in shPI4KIIα and shPI4KIIβ cells was a direct result of target protein depletion. The failure of GFP-tagged PI4KIIα-WT and PI4KIIβ-WT to fully restore TYRP1 localization might reflect partial interference with kinase activity by the C-terminal GFP fusion or a consequence of kinase overexpression.

**Figure 3.**
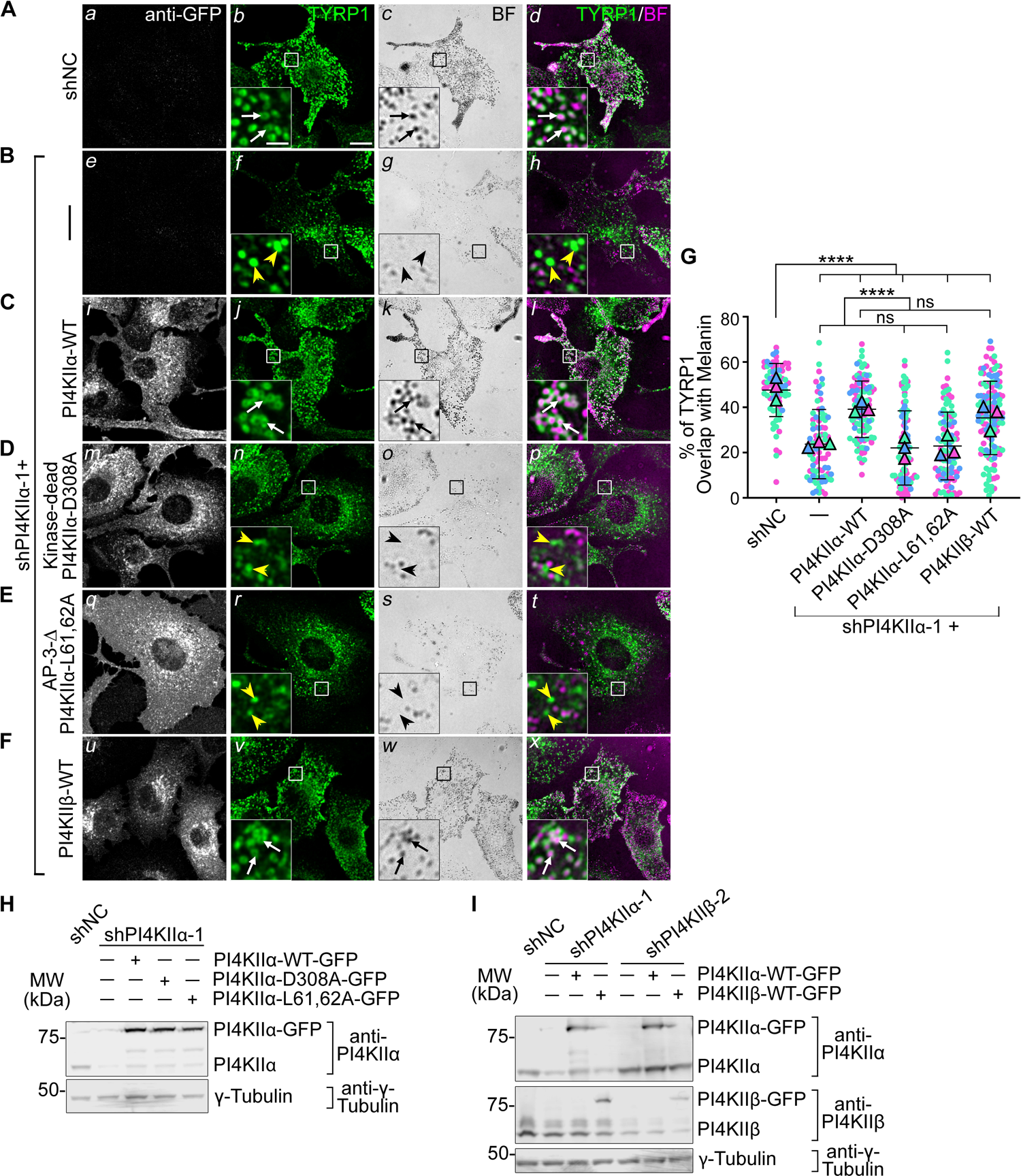
PI4KIIα function in TYRP1 trafficking requires its lipid kinase activity and binding to adaptor AP-3. Melan-Ink4a melanocytes were untransduced (A, B) or transduced with retroviruses to express GFP-tagged PI4KIIα-WT (C), kinase-dead mutant PI4KIIα-D308A (D), AP-3-binding-deficient mutant PI4KIIα-L61,62A (E) or PI4KIIβ-WT (F), selected with hygromycin for at least two weeks, and then transduced with lentiviruses to express shNC or shPI4KIIα-1. **(*A-F*)** 7 to 9 d later, cells were analyzed by IFM for GFP (white; left panels) and TYRP1 (green; middle left and right panels) and by bright field microscopy for pigment granules (BF; middle right panels; pseudocolored magenta at right). White arrows, TYRP1 surrounding pigment granules; yellow arrowheads, TYRP1 not overlapping with pigment granules. Boxed regions are magnified 5-fold in the insets. Scale bars: main panels, 10 μm; insets, 2 μm. **(*G*)** Quantification of TYRP1 overlap with melanosomes from at least three independent experiments, with analyses by Welch’s ANOVA; ****, p<0.0001; ns, not significant. **(*H*, *I*)** Whole-cell lysates from cells in A-E (H) or panel F and Fig. 4F (I) were analyzed by SDS/PAGE and immunoblotting for PI4KIIα (H) or both PI4KIIα and PI4KIIβ (I) and γ-Tubulin as a loading control. Left, positions of molecular weight markers (MW).

**Figure 4.**
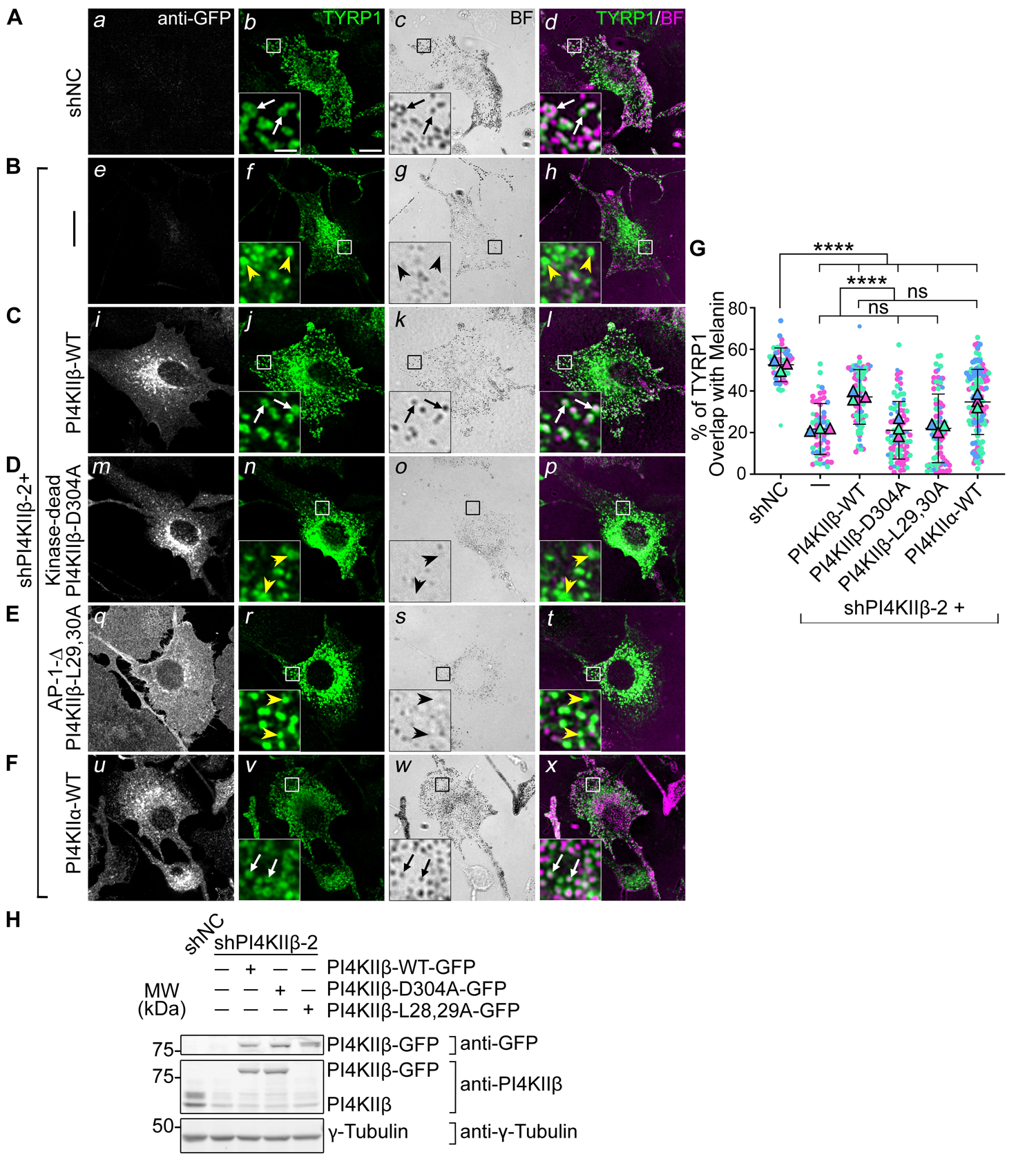
PI4KIIβ function in TYRP1 trafficking requires its lipid kinase activity and binding to adaptor AP-1. Melan-Ink4a cells were untransduced (A, B) or transduced with retroviruses to express GFP-tagged PI4KIIβ-WT (C), kinase-dead mutant PI4KIIβ-D304A (D), AP-1-binding-deficient mutant PI4KIIβ-L29,30A (E) or PI4KIIα-WT (F), selected for at least two weeks, and then transduced with lentiviruses expressing shNC or shPI4KIIβ-2 shRNA to PI4KIIβ. **(*A-F*)** 7 to 9 d after shRNA transduction, cells were analyzed by IFM for GFP (white; left panels) and TYRP1 (green; middle left and right panels) and by bright field microscopy for pigment granules (BF; middle right panels and pseudocolored magenta at right). White arrows, TYRP1 surrounding pigment granules; yellow arrowheads, TYRP1 not overlapping with pigment granules. Boxed regions are magnified 5-fold in the insets. Scale bars: main panels, 10 μm; insets, 2 μm. **(*G*)** Quantification of TYRP1 overlap with pigment granules from at least three independent experiments and analysis by Welch’s ANOVA; ****, p<0.0001; ns, not significant. **(*H*)** Whole-cell lysates from cells in A-E were analyzed by SDS/PAGE and immunoblotting for GFP, PI4KIIβ and γ-Tubulin as a loading control (note, the L28,29A mutant is not recognized by the anti-PI4KIIβ antibody). Left, positions of molecular weight markers (MW). Immunoblotting for PI4KIIα in cells (F) is shown in Figure 3.

Surprisingly, overexpression of either PI4KIIβ-WT-GFP in shPI4KIIα cells or of PI4KIIα-WT-GFP in shPI4KIIβ cells restored TYRP1 localization to the same degree as re-expression of the targeted kinase (white arrows, **Figs. 3F, G, I**; **Figs. 4F, G**). This likely reflects cross-compensation by overexpressed PtdIns-4-kinase activity in adjacent endosomal domains (see **Figs. 6-8** below).

We next tested if the PI4KIIα and PI4KIIβ roles in melanosome cargo transport require their kinase activities and respective binding to AP-3 or AP-1. Kinase-dead [PI4KIIα-D308A and PI4KIIβ-D304A; (Balla et al., 2002)] and AP binding-deficient mutants [PI4KIIα-L61, 62A and PI4KIIβ-L29, 30A; (Craige et al., 2008; Wieffer et al., 2013)] were expressed stably in WT mouse melanocytes, and the corresponding endogenous mouse kinase was depleted by shRNA for 7-9 d prior to analyses. IFM (**Fig. 3D, E**; **Fig. 4D, E**) and immunoblotting (**Figs. 3H, 4H**) showed that all mutants were effectively expressed. However, neither the kinase dead nor AP binding mutants of either PI4KIIα or PI4KIIβ restored TYRP1 localization to pigment granules in shPI4KIIα and shPI4KIIβ cells (yellow arrowheads; **Figs. 3G, 4G**). Moreover, whereas WT (**Figs. 3Ci, 4Ci**) and kinase-dead (**Fig. 3Dm, 4Dm**) variants of PI4KIIα-GFP and PI4KIIβ-GFP localized predominantly to cytoplasmic puncta (PI4KIIα) and to the perinuclear region or a few cytoplasmic puncta (PI4KIIβ), PI4KIIα-L61,62A-GFP (**Fig. 3Eq**) and PI4KIIβ-L29,30A-GFP (**Fig. 4Eq**) were predominantly cytoplasmic. These data suggest that (1) both PI4KIIα and PI4KIIβ require both their kinase activity (and hence PtdIns4P generation) and their interaction with APs to promote TYRP1 localization, and (2) recruitment to endolysosomal membranes requires binding of PI4KIIα to AP-3 and of PI4KIIβ to AP-1.

### PI4KIIα and PI4KIIβ independently function downstream of BLOC-1 in tubular transport to melanosomes

Because TYRP1 was trapped less in early endosomes upon PI4KIIα or PI4KIIβ depletion than upon BLOC-1 depletion/ knockout, we hypothesized that while either PI4KIIα or PI4KIIβ can collaborate with BLOC-1 to facilitate cargo exit from early sorting endosomes (Jani et al., 2021), each additionally promotes an ensuing step in tubule stabilization and/or targeting towards melanosomes. If so, then TYRP1 should remain trapped in early endosomes and not progress to late endosomes/ lysosomes upon PI4KIIα or PI4KIIβ depletion in BLOC-1-deficient cells. To test this, BLOC-1-deficient (BLOC-1^-/-^) melanocytes that lack the Pallidin (*Bloc1s6*) subunit and thus fail to assemble BLOC-1 (Falcon-Perez et al., 2002; Starcevic and Dell’Angelica, 2004) were treated with shNC, shPI4KIIα, or shPI4KIIβ. They were then analyzed by triple labeling IFM for TYRP1, either PI4KIIα or PI4KIIβ (not shown) to validate protein depletion, and transferrin receptor (TfR) to identify early sorting and recycling endosomes [TfR was used instead of STX13 to allow for triple labeling, and at steady state overlaps with STX13 but not TYRP1 in WT cells or with both STX13 and TYRP1 in BLOC-1^-/-^ cells (**Fig S4**)]. Consistent with published data (Delevoye et al., 2016; Setty et al., 2008; Setty et al., 2007) and **Fig. 2A**, TYRP1 in BLOC-1^-/-^ cells treated with shNC localized largely to early endosomes as shown by overlap with TfR (arrowheads, **Fig. 5Aa-c, 5B).** Knock down of PI4KIIα or PI4KIIβ in these cells with either of two shRNAs each did not alter the degree of TYRP1/ TfR overlap (yellow arrowheads, **Fig. 5Ad-i, 5B**). Thus, TYRP1 is trapped in early endosomes in BLOC-1-deficient cells regardless of the expression of PI4KIIα or PI4KIIβ, indicating that PI4KIIα and PI4KIIβ have unique functions downstream of BLOC-1 in cargo transport to melanosomes.

**Figure 5.**
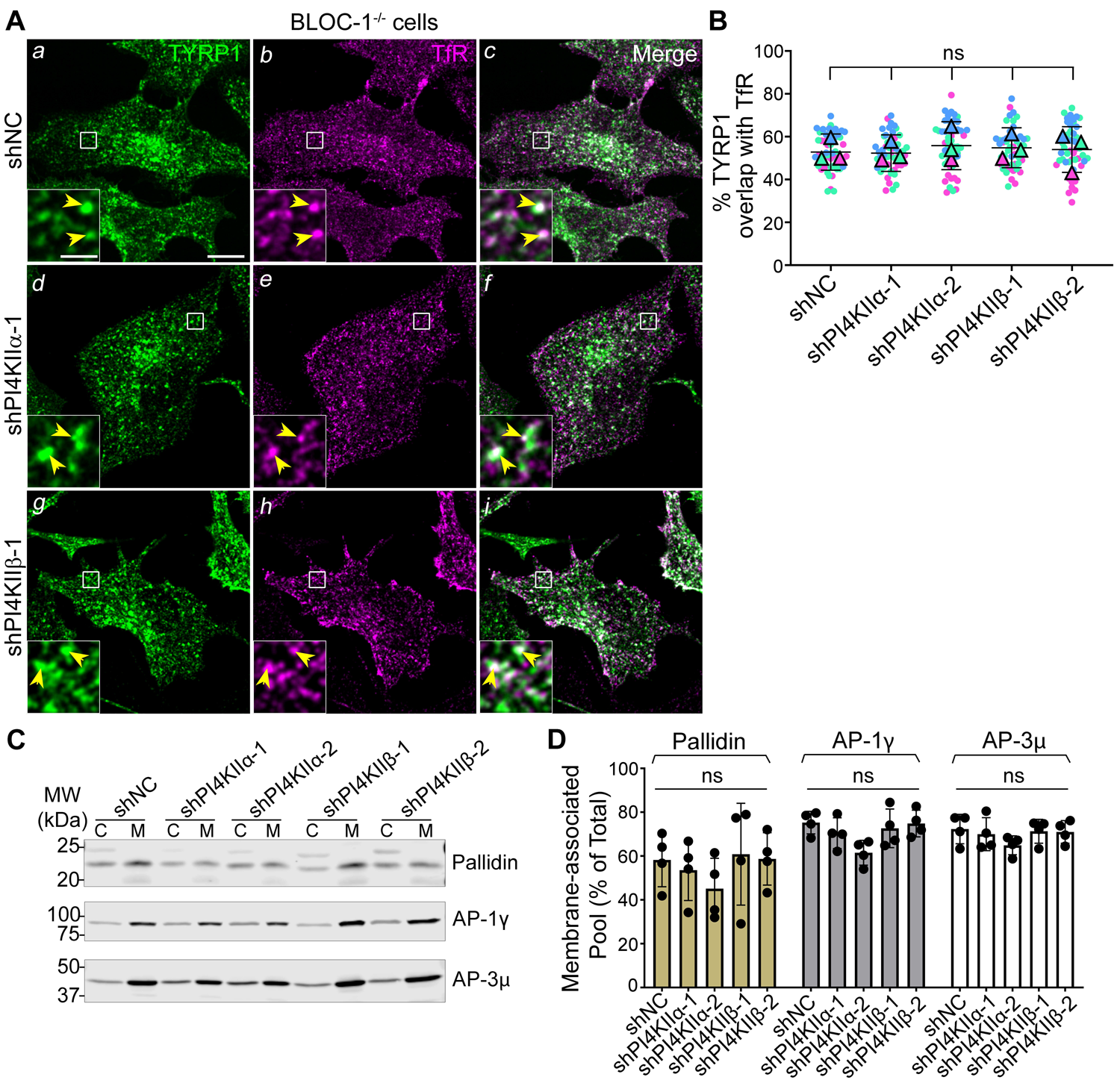
Type II PtdIns4-kinases are required after BLOC-1 function in tubular transport to melanosomes. BLOC-1^-/-^ melan-pa (A and B) or melan-Ink4a (C and D) melanocytes were transduced with lentiviruses to express shNC, shPI4KIIα, or shPI4KIIβ for 7 to 9 d. **(*A*)** BLOC-1^-/-^ cells were analyzed by IFM for TYRP1 (green; left and right panels), TfR to mark early endosomes (magenta, middle and right panels), and either PI4KIIα or PI4KIIβ (not shown); only cells with minimal PI4KIIα or PI4KIIβ signal are shown. Yellow arrowheads, TYRP1 overlapping with TfR. Boxed regions are magnified 5-fold in the insets. Scale bars: main panels, 10 μm; insets, 2 μm. (***B***) Quantification of TYRP1 overlap with TfR from three independent experiments and analysis by ordinary one-way ANOVA. **(*C*, *D*)** Indicated melan-Ink4a cells were homogenized and fractionated to yield postnuclear membrane (M) and cytosolic (C) fractions. Identical cell equivalents of membrane and cytosolic fractions were fractionated by SDS-PAGE and analyzed by immunoblotting using antibodies to the indicated subunits of BLOC-1, AP-1, and AP-3. **(*C*)** Representative blots. Left, positions of molecular weight (MW) markers. **(*D*)** Quantification of the percentage (mean ± SD of 4 independent experiments) of each protein associated with the membrane fraction relative to the total cellular content. Analysis, ordinary one-way ANOVA. ns, not significant.

PI4KIIα and PI4KIIβ could potentially generate a pool of PtdIns4P to recruit or stabilize BLOC-1 or other effectors of tubular transport on membranes, as suggested by *in vitro* studies (Jani et al., 2021). Because validated fluorescent forms of BLOC-1 do not exist to assess membrane association by live cell microscopy in which intact tubules can be visualized, we instead used subcellular fractionation. Membrane and cytosolic fractions were prepared from homogenates of WT melanocytes treated with shNC, shPI4KIIα, or shPI4KIIβ, and the relative association of BLOC-1, AP-1, and AP-3 with each fraction was assessed by immunoblotting for the Pallidin, AP-1γ, and AP-3μ subunits, respectively (**Fig. 5C, D**). Depletion of PI4KIIα or PI4KIIβ neither substantially nor significantly reduced BLOC-1 membrane association (**Fig. 5C, D**; the decrease caused by shPI4KIIα-2 likely reflects an off-target effect as discussed earlier). Similarly, neither shPI4KIIα nor shPI4KIIβ affected the membrane association of AP-1 or AP-3 (**Fig. 5C, D**). We therefore conclude that neither type II PtdIns4-kinase is uniquely required for the association of BLOC-1, AP-3 or AP-1 with membranes.

### PI4KIIα and PI4KIIβ function sequentially to enrich PtdIns4P on tubular endosomal domains

If PI4KIIα and PI4KIIβ each directly regulate cargo transport downstream of BLOC-1, then they and their product, PtdIns4P, should be present on early endosomes and the tubular carriers through which BLOC-1-dependent cargoes travel. By IFM analysis of fixed cells, endogenous PI4KIIα and PI4KIIβ and overexpressed GFP-tagged PI4KIIα and PI4KIIβ localized in the cell periphery to only a small fraction of early endosomes labeled by TfR (yellow arrowheads, **Fig. 6A, B**) or STX13 (yellow arrowheads, **Fig. 6C, D**), respectively (note also extensive localization of PI4KIIβ to the Golgi region). Similarly small fractions of endogenous PI4KIIα and PI4KIIβ colocalized with TfR [as seen in other cell types (Craige et al., 2008; Wieffer et al., 2013), but a larger fraction of GFP-tagged forms colocalized with STX13 – suggesting that PI4KIIα and PI4KIIβ in the cell periphery both predominantly localize to a small subset of early endosomal structures that are more enriched in STX13 than TfR.

**Figure 6.**
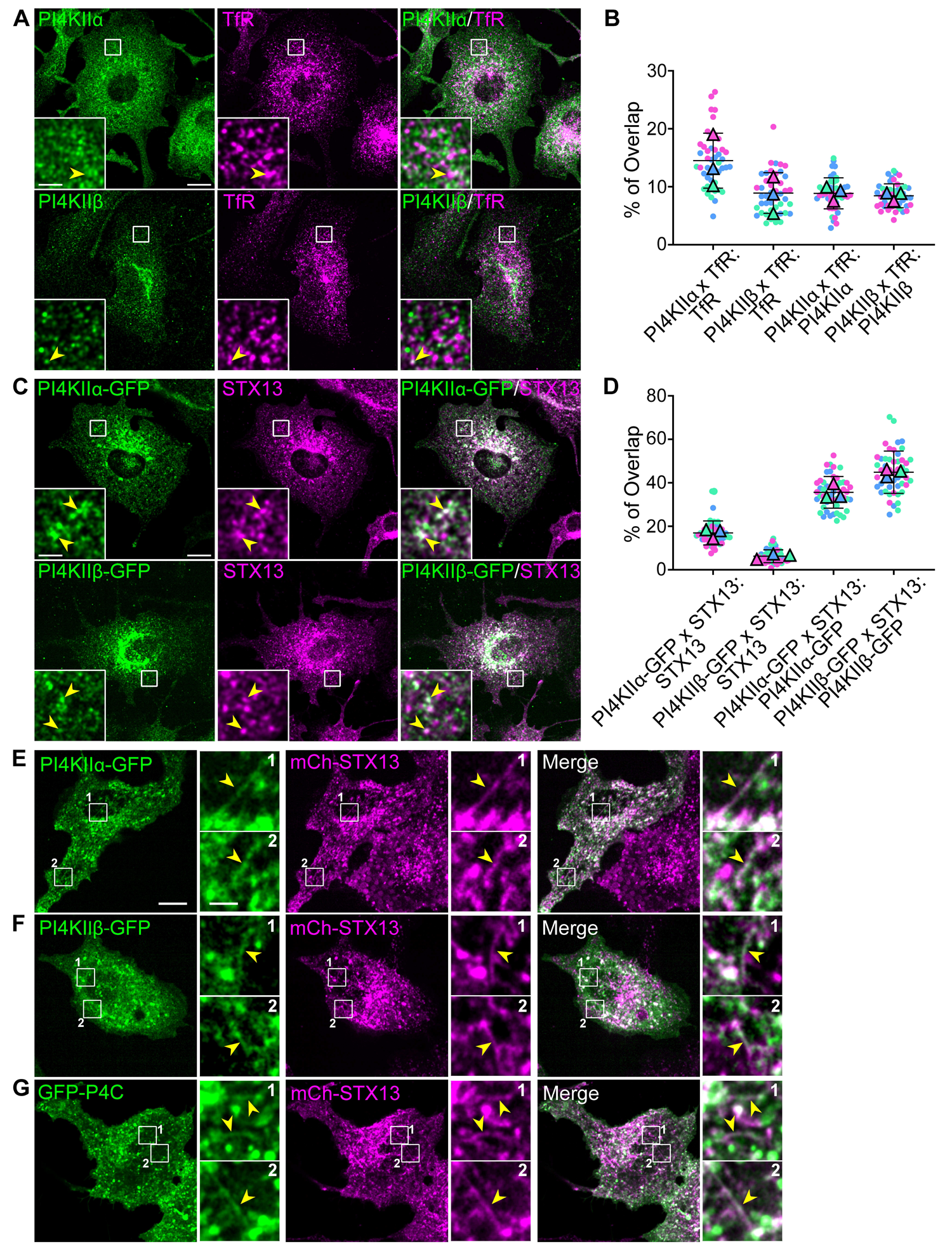
Type II PtdIns4-kinases and PtdIns4P are present on endosomal tubular carriers. (*A-D*) Melan-Ink4a cells that were untransfected (A, B) or that stably expressed PI4KIIα-GFP or PI4KIIβ-GFP (C, D) were fixed and analyzed by IFM for either endogenous TfR with PI4KIIα or PI4KIIβ (A, B) or endogenous STX13 with GFP-tagged PI4KIIα or PI4KIIβ (*C*, *D*). Shown are representative individual and merged images (A, C) and quantification of the percentage of area of overlap of both contents (mean ± SD from three experiments) relative to the total area labeled by either protein alone (B, D). **(*E*-*G*)** Melan-Ink4a cells stably expressing mCherry-STX13 (magenta, middle and right panels) were transiently transfected with GFP-tagged PI4KIIα (E), PI4KIIβ (F) or P4C (G; green, left and right panels) and then analyzed by dual-view live cell-spinning disk microscopy. Shown are the individual labels and merged image from a frame of a representative cell emphasizing tubular structures; two boxed regions are magnified 5-fold in the insets. Yellow arrowheads, tubular structures containing both mCherry-STX13 and either PI4KIIα-GFP, PI4KIIβ-GFP, or GFP-P4C-SidC. Scale bars: main panels, 10 μm; insets, 2 μm.

To better assess localization to the fixation-sensitive tubules, we utilized live-cell microscopy to visualize GFP-tagged PI4KIIα, PI4KIIβ or a PtdIns4P probe relative to mCherry-tagged STX13, which labels the entire early endosomal network including the tubular carriers (Bowman et al., 2021; Delevoye et al., 2016; Dennis et al., 2015). To detect PtdIns4P, we used P4C-SidC (P4C), a PtdIns4P-binding domain derived from the secreted SidC effector protein of *Legionella pneumophila* (Dolinsky et al., 2014; Weber et al., 2018; Weber et al., 2014) [GFP-P4M-SidMx2 – containing a tandem repeat of the PtdIns4P binding domain of SidM from *L. pneumophila* (Hammond et al., 2014) – did not efficiently label Golgi, endosomes, or tubules in our cells, and thus was not used in our analyses]. GFP-tagged proteins were transiently expressed in WT melanocytes stably expressing mCherry-STX13, and cells were analyzed the next day by dual-view live-cell spinning disk microscopy (Kinosita et al., 1991) to simultaneously detect mCherry- and GFP-tagged components. Still images from image streams show that GFP-tagged PI4KIIα, PI4KIIβ, and P4C each labeled punctate structures and tubules that largely overlapped with mCherry-STX13 (yellow arrowheads in insets, **Fig. 6A-C**), consistent with previous studies (Balla et al., 2002; Craige et al., 2008; Dong et al., 2016; Jovic et al., 2014; Ma et al., 2020; Wieffer et al., 2013). Since the majority of STX13-labeled tubules in melanocytes are BLOC-1-dependent (Delevoye et al., 2016), the tubules decorated by both mCherry-STX13 and PI4KIIα-GFP, PI4KIIβ-GFP, or GFP-P4C likely corresponded to the BLOC-1-dependent tubular carriers. Additional tubules were labeled only by PI4KIIα-GFP, PI4KIIβ-GFP, or GFP-P4C (data not shown), suggesting that type II PtdIns4-kinases associate with tubules from multiple origins in melanocytes. As expected, P4C also labeled the plasma membrane and the Golgi area (**Fig. 6C**; see also **Fig. 8Aa-c**). The data suggest that PI4KIIα, PI4KIIβ, and their product PtdIns4P are present on early endosomal tubules that ferry cargo to melanosomes and on other tubules.

Since both PI4KIIα and PI4KIIβ associated with membrane tubules, we asked whether the requirement for both in cargo delivery to melanosomes reflects their sequential activity. To test this, we analyzed the dual-view image streams for the temporal association of PI4KIIα-GFP or PI4KIIβ-GFP relative to the onset of tubule formation from round STX13-containing early endosomes. For image streams in which the source endosome and a nascent tubule were visualized, PI4KIIα-GFP was consistently present both on the body of mCherry-STX13-containing endosomes prior to tubule formation (**Fig. 7A**, white arrowheads) and on the nascent mCherry-STX13-labeled tubule, starting with the appearance of a small protrusion (**Fig. 7A**, yellow arrowhead, 0s) and continuing during tubule extension (**Fig. 7A**, yellow arrowheads, 0.4-4s). This indicated that PI4KIIα is sorted into the tubular carriers during their formation. By contrast, PI4KIIβ-GFP was not detected on the nascent mCherry-STX13-labeled tubules (**Fig. 7B**; tubule initiates at white arrowhead), but rather consistently appeared as the tubule extended and accumulated along its length (**Fig. 7B**, 1.6-2.4s between white and yellow arrowheads). In cases where PI4KIIβ-GFP was detected on the source endosome prior to tubulation, it was typically present in a subdomain (**Fig. 7B**, white arrowheads) and became more enriched on the tubules. These results indicate that PI4KIIβ, unlike PI4KIIα, associates with established tubules, and suggest that PI4KIIβ functions at a later step of tubule dynamics than PI4KIIα.

**Figure 7.**
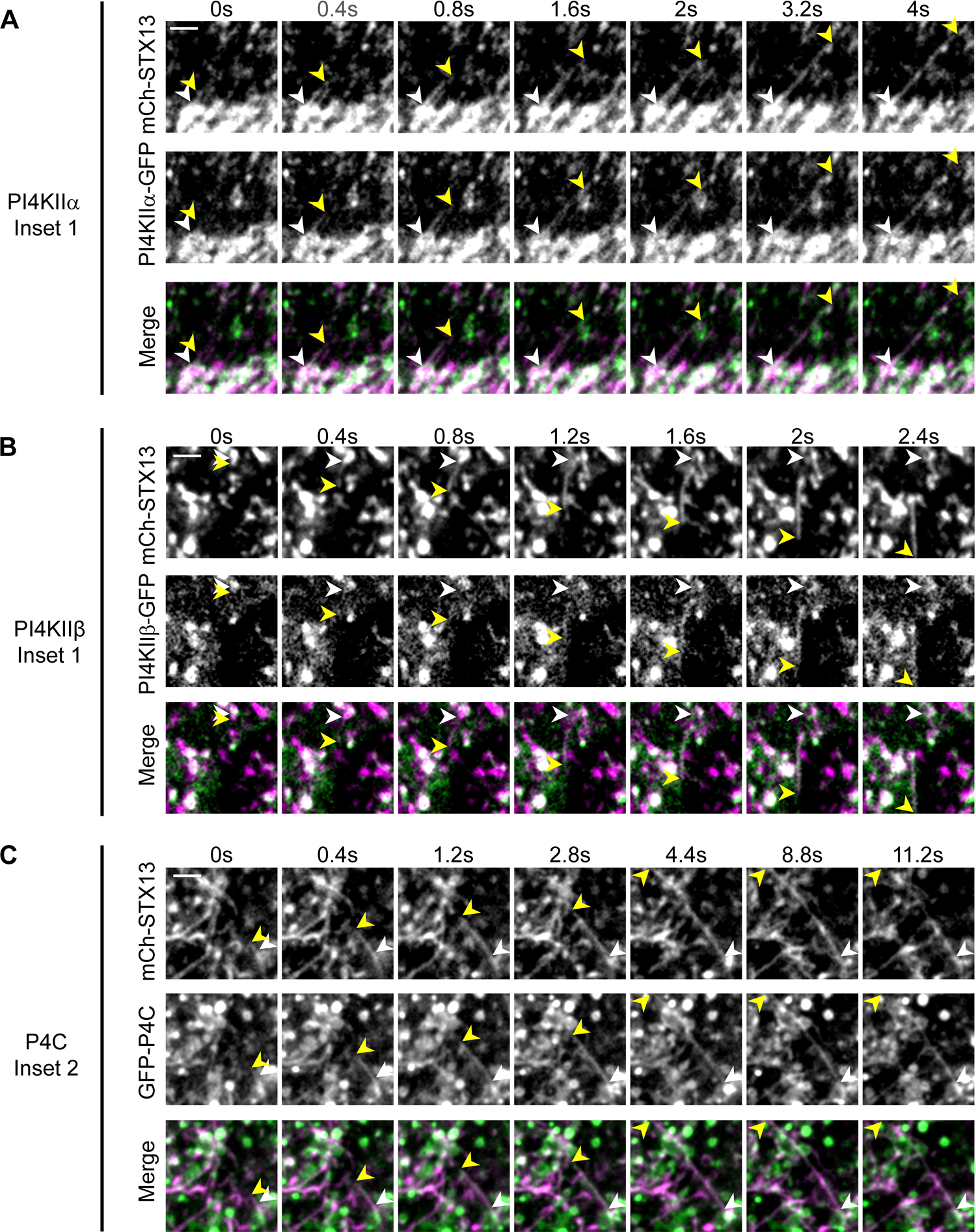
PI4KIIα and PI4KIIβ associate with tubules with distinct kinetics and together generate PtdIns4P throughout the lifetime of the tubule. As in Fig. 6E-G, melan-Ink4a cells stably expressing mCherry-STX13 and transiently transfected to express GFP-tagged PI4KIIα (A), PI4KIIβ (B) or P4C (C) were analyzed by dual-view live-cell spinning-disk microscopy. Shown are a montage of individual and merged image sequences from the indicated insets from Fig. 6E-G emphasizing the successive extension of STX13-containing tubules from a source endosome. White arrowheads, sources of tubules; yellow arrowheads, tips of tubules; time from onset of tubule formation is indicated in seconds. GFP-PI4KIIα was present on nascent tubules in 22 out of 24 instances (as in A; n=6 cells, 3 independent experiments) and GFP-PI4KIIβ joined pre-formed tubules in 20 out of 26 instances (as in B, n=9 cells, 3 independent experiments) in which the source endosome and a nascent tubule were visible and distinguishable in the same plane of focus. Scale bars: 2 μm.

Consistent with the combined distribution of PI4KIIα-GFP and PI4KIIβ-GFP on the tubules, their product, PtdIns4P (visualized with GFP-P4C), was present throughout the lifetime of the mCherry-STX13-positve tubules from the onset of tubule formation through tubule extension (**Fig. 7C**, all panels from white arrows to yellow arrowheads). Additional tubules labeled by GFP-P4C but not mCherry-STX13 were observed moving towards the plasma membrane and likely represent Golgi-derived carriers (Graham and Burd, 2011; Highland and Fromme, 2021; Stalder and Gershlick, 2020).

### PI4KIIα and PI4KIIβ are both required to maintain the endosomal PtdIns4P pool

To test whether PI4KIIα and PI4KIIβ activity was responsible for PtdIns4P on vacuolar endosomes and endosomal tubules, we assessed the distribution of PtdIns4P in shPI4KIIα and shPI4KIIβ cells using GFP-P4C. To avoid the heterogeneity of transient expression, GFP-P4C was expressed stably from a recombinant retrovirus in WT melanocytes prior to expression of shNC, shPI4KIIα, or shPI4KIIβ from recombinant lentiviruses. Cells were then transiently transfected to express mCherry-STX13 prior to live-cell microscopy. As in transient transfections (**Fig. 6**), GFP-P4C expressed stably in shNC cells localized to the plasma membrane, Golgi, and perinuclear and peripheral tubular and punctate compartments (**Fig. 8Aa**). Depletion of PI4KIIα or PI4KIIβ resulted in an apparent diminution of GFP-P4C-labeled puncta in the periphery and an accumulation in the perinuclear region (**Fig. 8Ad, g**); unlike in cells that did not express P4C (**Fig. 2A**), this was accompanied by perinuclear accumulation of STX13-labeled endosomes (**Fig. 8Ae, h**), perhaps due to exacerbation of altered endosomal dynamics by shielding of remaining PtdIns4P. Concomitantly, compared to shNC cells, fewer tubular structures that were labeled by GFP-P4C (**Fig. 8Aa,d, g**; **Fig. 8B**) or doubly labeled by both GFP-P4C and mCherry-STX13 (yellow vs. white arrowheads; **Fig. 8Ac, f, i**; **Fig. 8C**) were observed in shPI4KIIα and shPI4KIIβ cells. Moreover, the percentage of mCherry-STX13-containing tubules that also labeled for GFP-P4C was dramatically reduced (**Fig. 8D**). The impact of PI4KIIα- and PI4KIIβ-depletion was specific for PtdIns4P, since the number of puncta and the general distribution of the PtdIns3P probe GFP-2XFYVE (Gillooly et al., 2000) were similar in cells expressing shNC, shPI4KIIα, or shPI4KIIβ (**Fig. S5**). These data show that both PI4KIIα and PI4KIIβ contribute to PtdIns4P pools on membrane tubules emanating from early endosomes and other compartments, such as lysosomes (Levin-Konigsberg et al., 2019).

**Figure 8.**
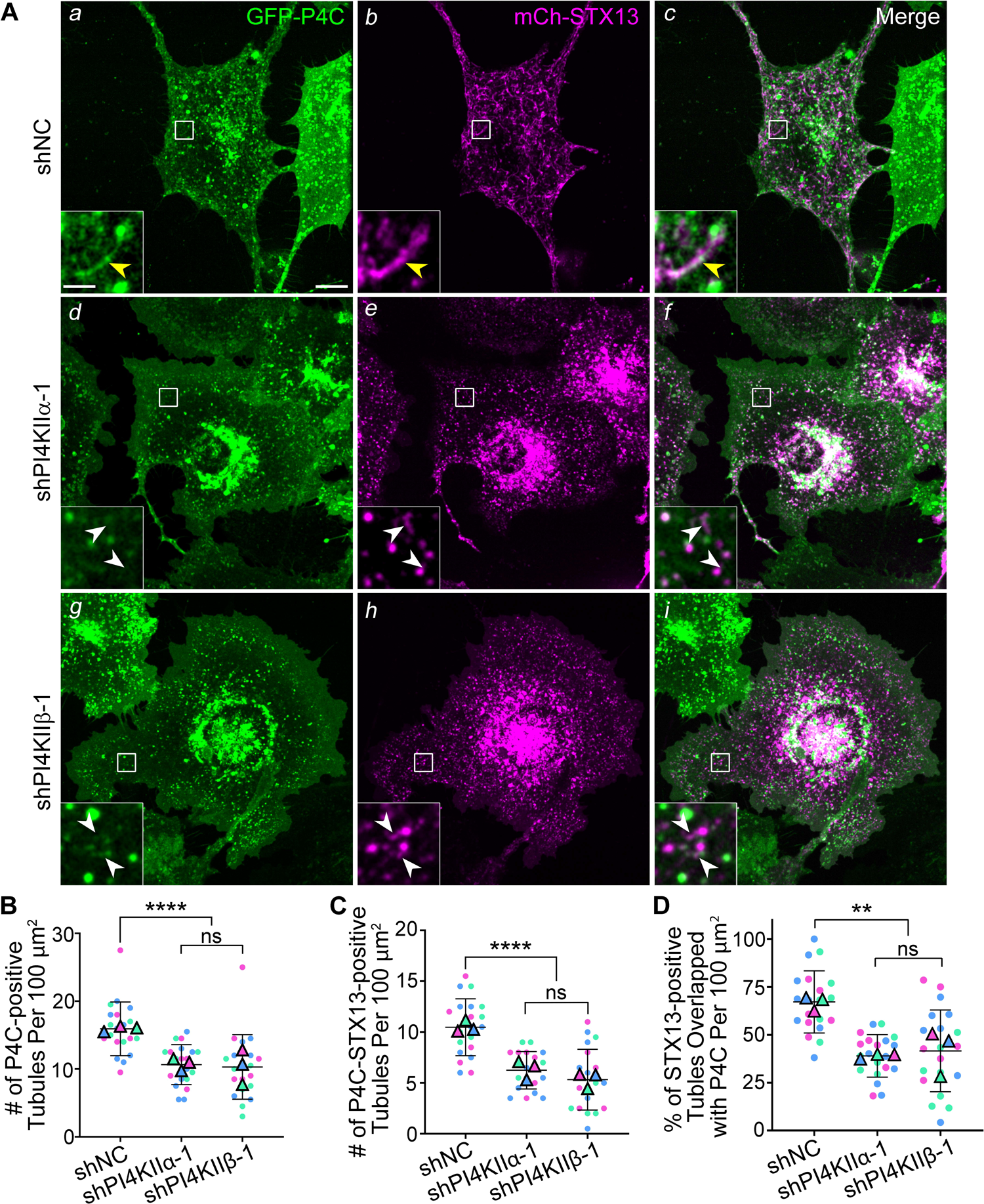
PI4KIIα and PI4KIIβ are both required to maintain PtdIns4P on endosomal tubules. Melan-Ink4a cells stably expressing GFP-P4C (P4C) were transduced with lentiviruses encoding shNC or the indicated shRNAs to PI4KIIα or PI4KIIβ, selected for 7 to 9 d, and then transiently transfected with mCherry-STX13 (STX13) and analyzed by live-cell spinning-disk microscopy. **(*A*)** Still images from image streams of cells treated with shNC (Aa-c) shPI4KIIα (Ad-f) or shPI4KIIβ (Ag-i) showing P4C (green, left panels), STX13 (magenta, middle panels), and their overlap (merged images on the right). Yellow arrowheads, STX13-containing tubules that overlap with P4C; white arrowheads, STX13-containing tubules that do not overlap with P4C. Boxed regions are magnified 5-fold in the insets. Scale bars: main panels, 10 μm; insets, 2 μm. **(*B*, *C*)** Quantification of the total numbers of either single P4C-positive (B) or double P4C- and STX13-positive tubules per 100-μm^2^ area in each cell population. **(*D*)** Quantification of the percentage of STX13-containing tubules that were also positive for P4C in each cell population. Data in B-D are from three independent experiments each and were analysed by Kruskal-Wallis (B) or ordinary one-way ANOVA (C, D). ****, p<0.0001; **, p<0.01; ns, not significant.

### PI4KIIα and PI4KIIβ are needed to support the formation of effective tubular carriers

Given that PI4KIIα and PI4KIIβ localization correlated with tubule formation and elongation respectively, we next tested whether PtdIns4P production by PI4KIIα and PI4KIIβ is required for proper tubule dynamics in melanocytes. WT melanocytes that stably expressed mCherry-STX13 were treated with shNC, shPI4KIIα, or shPI4KIIβ, and the effects on STX13-positive tubular structures were assessed by live-cell microscopy. In shNC cells, as in untreated cells [**Figs. 6** and **7** and (Bowman et al., 2021; Dennis et al., 2016; Dennis et al., 2015)], numerous long and dynamic STX13-positive tubules were detected throughout the cytoplasm [(shown are fluorescence (**Fig 9Aa, b**) and skeletonized images (**Fig. 9Ac, d**)]. Although shPI4KIIα and shPI4KIIβ cells also harbored dynamic tubules, relative to shNC cells the tubules were fewer in number (**Fig. 9Aa-d, Ba-d, Ca-d**; **Fig. 9D**), less enriched in the number and percentage of longer (> 1 μm) tubules (**Fig. 9E, 9F**), and shorter on average (**Fig. 9G**). Since the tubules that contact melanosomes are typically long (Bowman et al., 2021; Dennis et al., 2015), these data suggest that formation and extension of melanosome-bound tubules are impaired by PI4KIIα or PI4KIIβ deficiency. The reduction in tubule number and length trended as more severe in shPI4KIIα cells than in shPI4KIIβ cells (**Fig. 9D-G**), supporting the conclusion that PI4KIIα functions earlier than PI4KIIβ, but were less severe than previously observed in BLOC-1^-/-^ cells (Bowman et al., 2021; Delevoye et al., 2016). These results indicate that PI4KIIα and PI4KIIβ are each uniquely required sequentially in tubule elongation and/or stabilization, subsequent to BLOC-1-dependent tubule formation.

**Figure 9.**
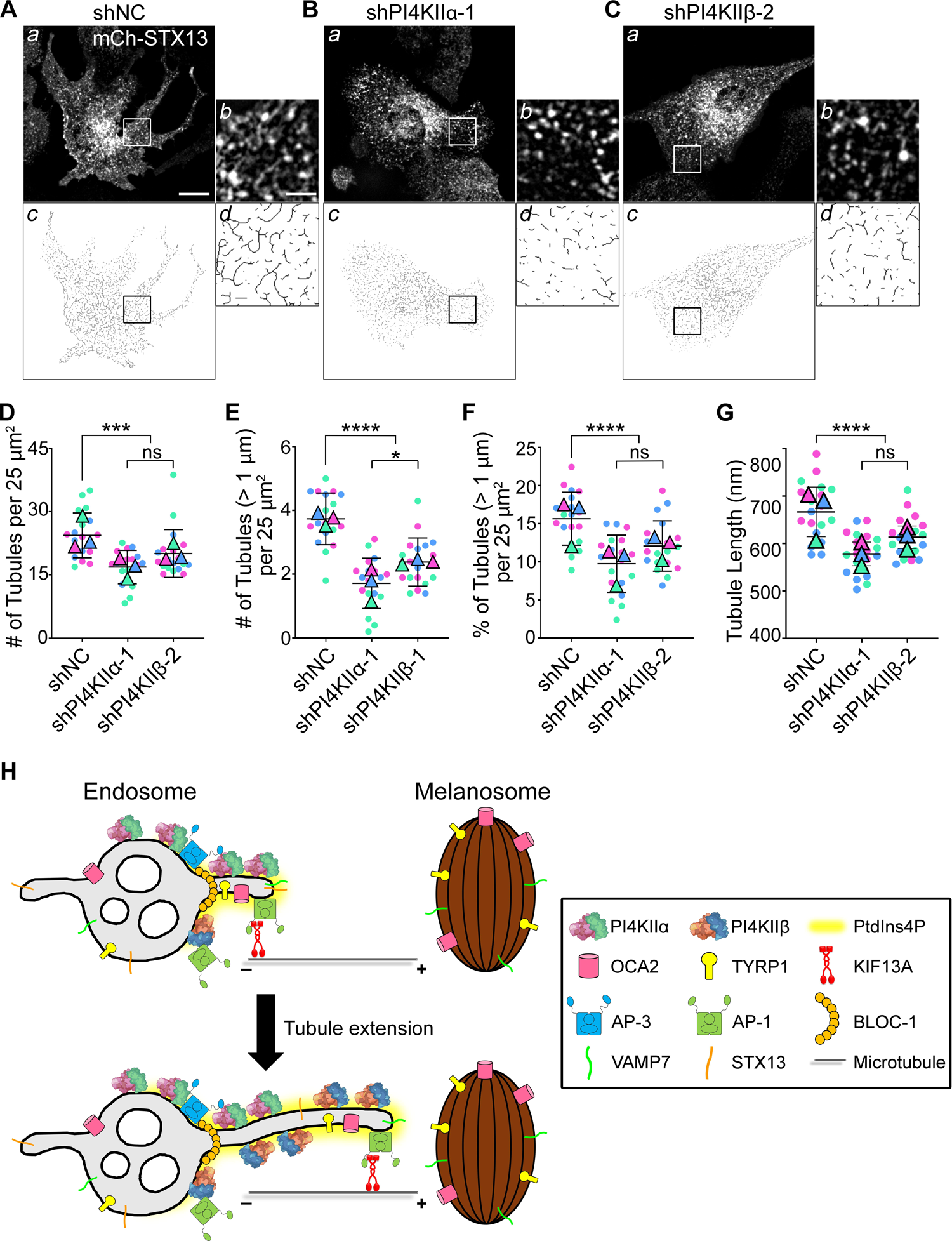
Depletion of either PI4KIIα or PI4KIIβ disrupts the dynamics of BLOC-1-dependent tubular cargo carriers. ***A-G.*** Melan-Ink4a melanocytes stably expressing mCherry-STX13 (mCh-STX13) were transduced with lentiviruses to express shNC or the indicated shRNAs to PI4KIIα or PI4KIIβ, selected for 7 to 9 d, and analyzed by live cell-spinning disk microscopy. **(*A*-*C*)** Still images from image streams (a panels; boxed regions are magnified 4-fold in insets in b panels), and skeletonized images (c, d) emphasizing the tubules. Scale bars: main panels, 12 μm; insets, 3 μm. **(*D****-**G***) Quantification of the total number of mCh-STX13-labeled tubules per 25-μm^2^ area (D), the total number of tubules per 25-μm^2^ area that were longer than 1 μm (E), the percentage of total tubules that were longer than 1 μm (F), or the length of all tubules per 25-μm^2^ area (G) for each cell population. Data are from three independent experiments and were analyzed by Kruskal-Wallis (D) or ordinary one-way ANOVA (E-G). (****, p<0.0001; ***, p<0.001; *, p<0.05; NS, not significant). **(*H*)** Working model. PI4KIIα (pink and green) is recruited by AP-3 (blue) on the vacuolar endosome to newly forming endosomal tubules labeled by STX13 through the interaction of AP-3, BLOC-1 (orange), and PI4KIIα (top panel). As the tubule elongates, PI4KIIβ (blue and green) is recruited by AP-1 (green) – perhaps bound to the kinesin motor heavy chain KIF13A (red) – to the extended tubule (bottom panel). Both PI4KIIα and PI4KIIβ generate local PtdIns4P (yellow shading) to allow for effective transport of cargos (TYRP1, OCA2 and VAMP7 as indicated) to melanosomes.

## DISCUSSION

BLOC-1-dependent endosomal tubular carriers are essential for cargo delivery to maturing melanosomes, but how lipids regulate tubule formation and dynamics was not known. We show that the type II PtdIns-4-kinases, PI4KIIα and PI4KIIβ, are recruited to the tubular endosomal carriers where they cooperate to generate local pools of PtdIns4P. While PI4KIIα and PI4KIIβ likely play redundant roles in PtdIns4P generation on vacuolar endosomes to concentrate BLOC-1 and initiate tubulation (Jani et al., 2021), the PtdIns4P pools produced by each kinase are required after tubule initiation for tubule stability and ultimate cargo delivery to melanosomes. These functions require both kinase activity and adaptor binding by PI4KIIα and PI4KIIβ. Depletion of either PI4KIIα or PI4KIIβ impairs tubular carrier dynamics, resulting in melanosomal cargo mistargeting – primarily to late endosomes and lysosomes – and consequent impaired melanin production. Our results support a model in which PI4KIIα on newly generated endosomal tubules and PI4KIIβ on the growing tubules, recruited by AP-3 and AP-1 respectively, generate local PtdIns4P to ensure effective tubular cargo transport to melanosomes (**Fig. 9H**). This model may have broader implications for tubule-dependent cargo transport within the endomembrane system and for LRO biogenesis in general.

In melanocytes, BLOC-1 and the kinesin-3 heavy chain KIF13A cooperate to generate long, stable membrane tubules that transport selected cargoes to melanosomes and constitute the majority of STX13-containing endosomal tubules (Delevoye et al., 2016; Dennis et al., 2016; Dennis et al., 2015). BLOC-1 cooperates with PI4KIIα in endosomal trafficking in other cell types (Gokhale et al., 2012; Larimore et al., 2011; Salazar et al., 2009), but how PI4KIIα or PI4KIIb contributes to tubule dynamics was not known. Our live-cell imaging shows that PtdIns4P is present over the entire length and lifetime of BLOC-1-dependent tubules. PI4KIIα and PI4KIIβ are both detected on the tubules and the early endosomes from which they emerge, and accordingly, depletion of either PI4KIIα or PI4KIIβ impaired both tubule number and length. Thus, continuous PtdIns4P generation by PI4KIIα and PI4KIIβ coincides with and is necessary for tubule initiation, extension, and stabilization. PI4KIIα or PI4KIIβ depletion impaired tubule number less dramatically than BLOC-1 depletion; we interpret this to reflect redundancy of the two type II PtdIns-4-kinases in tubule initiation (Jani et al., 2021). Our data in melanocytes are consistent with the known association of PtdIns4P with recycling endosomal tubules in HeLa cells (Jovic et al., 2009) and with tubules from a variety of LROs in different cell types (Domingues et al., 2016; Levin-Konigsberg et al., 2019; López-Haber et al., 2020; Ma et al., 2020). How PtdIns4P promotes tubule stability or extension is not clear. We speculate that it serves as a platform to recruit membrane bending proteins (Kitamata et al., 2020) such as BLOC-1 itself [(Jani et al., 2021); see below], EHD1, or DRG2 (Jovic et al., 2009; Ma et al., 2020; Mani et al., 2017). Alternatively, PtdIns4P might concentrate KIF13A on tubules either indirectly via clustering its adaptor AP-1 (Wang et al., 2003) or directly like the activity of PtdIns(4,5)P_2_ on KIF1A and KIF5B to drive membrane transport and tubulation (Du et al., 2016; Klopfenstein et al., 2002).

Perhaps our most surprising finding is that two type II PtdIns-4-kinases are sequentially required for extension and/or stability of the same tubular transport carriers. Depletion of either PI4KIIα or PI4KIIβ reduced the number and length of STX13-positive tubules and the delivery of melanosome cargo. As in other cell types (Balla et al., 2002; Craige et al., 2008; Jovic et al., 2014; Ma et al., 2020; Wieffer et al., 2013), both PI4KIIα and PI4KIIβ in melanocytes localized in part to a subset of early endosomes and derived tubules, but whereas PI4KIIα associated with the tubules during their formation (and thus might play a prominent role in tubule initiation), PI4KIIβ mainly accumulated on the tubules after formation and during extension. Consistently, tubules were shorter upon depletion of PI4KIIα than of PI4KIIβ. Together, these data suggest that PI4KIIα and PI4KIIβ act sequentially to ensure that PtdIns4P levels are maintained throughout tubule elongation (**Fig. 9H**). Because PI4KIIα function and localization required its AP-3-binding determinant, we speculate that PI4KIIα is recruited by AP-3 on budding regions of vacuolar endosomes, where a BLOC-1-AP-3 super-complex (Di Pietro et al., 2006; Salazar et al., 2006; Salazar et al., 2009) facilitates tubule initiation and SNARE and cargo sorting (Bowman et al., 2021). Similarly, PI4KIIβ function and localization required its AP-1-binding determinant; we thus further speculate that PI4KIIβ is recruited to the growing tubule by AP-1 bound to KIF13A as KIF13A extends the tubule along microtubules (Campagne et al., 2018; Delevoye et al., 2016; Delevoye et al., 2009; Delevoye et al., 2014). Although general membrane association of BLOC-1, AP-3, or AP-1 was not impaired by PI4KIIα- or PI4KIIβ-depletion, the pool of PtdIns4P generated by PI4KIIα or PI4KIIβ likely feeds back to concentrate BLOC-1 (Jani et al., 2021) and AP-1 (Wang et al., 2003), respectively, on the tubules. Although overexpression of GFP-PI4KIIα or -PI4KIIβ could compensate for the loss of either endogenous PI4KII paralogue in cargo localization to melanosomes, excess local PtdIns4P produced by the overexpressed PI4KII likely diffuses to sites where the absent PI4KII paralogue would normally produce a more limited pool.

While our live cell imaging data are consistent with a redundant requirement for either type II PtdIns-4-kinase to concentrate BLOC-1 during recycling tubule formation from early endosomal vacuoles (Jani et al., 2021), they reveal an independent non-redundant requirement for PI4KIIα and PI4KIIβ in tubule dynamics and melanosome cargo transport downstream of BLOC-1. This is supported by: (a) the exit of BLOC-1-dependent melanosome cargoes from early endosomes in cells depleted of PI4KIIα or PI4KIIβ but not in cells depleted of BLOC-1; (b) the entrapment of cargo in early endosomes in cells lacking both BLOC-1 and either PI4KIIα or PI4KIIβ; (c) the lack of a significant reduction in BLOC-1 recruitment to membranes in cells depleted of PI4KIIα or PI4KIIβ; and (d) the more modest impairment in tubule number and length in cells depleted of PI4KIIα or PI4KIIβ relative to BLOC-1-deficient cells (Bowman et al., 2021; Delevoye et al., 2016). The additional requirement for PtdIns4P in the maintenance and extension of the tubules at a stage following BLOC-1-dependent tubule initiation is also consistent with data in HeLa cells (Jani et al., 2021). Our efforts to simultaneously deplete both enzymes in melanocytes were unsuccessful, but future studies using knockout/ knockdown approaches might be used to validate a requirement for endosomal PtdIns4P in initiating tubule formation in melanocytes.

Both type II PtdIns-4-kinases were necessary for the ultimate delivery of BLOC-1-dependent cargoes to melanosomes; depletion of either PI4KIIα or PI4KIIβ resulted in mistargeting to late endosomes and lysosomes for at least one cargo. This does not likely reflect improper sorting of cargo or SNAREs into the nascent tubules, as sorting is mediated by AP-1, AP-3, and/or BLOC-1 (Bowman et al., 2021; Delevoye et al., 2009; Sitaram et al., 2012; Theos et al., 2005), which were effectively recruited to membranes in PI4KIIα- or PI4KIIβ-depleted cells. Moreover, cargo was retained within early endosomes in BLOC-1-deficient cells depleted of PI4KIIα or PI4KIIb, indicating that missorting to late endosomes/ lysosomes required BLOC-1 activity and thus likely also tubule initiation. Rather, the data support the conclusion that the short tubules in PI4KIIα or PI4KIIb-depleted cells are sufficient for cargo exit from early endosomal vacuoles but not for ultimate cargo delivery to melanosomes. Thus, length and lifetime likely impact the ability of tubules to fuse with maturing melanosomes to ensure cargo delivery (Delevoye et al., 2009). This might reflect (a) the long distance needed for cargo to travel from endosomes to peripheral melanosomes or (b) a requirement for selective microtubule-driven processes to ensure that the cargo carriers encounter and fuse with melanosomes prior to late endosomes or lysosomes. Future studies will be needed to address this model.

PtdIns4P is critical in all cells to control cargo transport, membrane identity, and organelle contacts within the endomembrane system (D’Angelo et al., 2008; Tan and Brill, 2014). Specialized cell types specifically adapt the use of PtdIns4P on tubules to ensure the maturation of LROs, including phagolysosomes in dendritic cells and macrophages (Levin-Konigsberg et al., 2019; López-Haber et al., 2020; Mantegazza et al., 2014) and secretory granules in *Drosophila melanogaster* salivary glands (Ma et al., 2020). Our results extend these studies by revealing a requirement in melanocytes for both mammalian type II PtdIns-4-kinases at distinct stages of membrane tubule-dependent anterograde cargo transport from endosomes to maturing melanosomes, and are consistent with a requirement for both PI4KIIα and PI4KIIb in Weibel-Palade body maturation in endothelial cells (Lopes da Silva et al., 2016). Together, these studies suggest that the formation of PtdIns4P-enriched tubules is a general feature in LRO maturation.

## MATERIALS AND METHODS

### Reagents

Unless otherwise indicated, chemicals were obtained from Sigma-Aldrich or Fisher Bioreagents. Tissue culture reagents and blasticidin were from Life Technologies/Thermo Fisher Scientific. Puromycin was from Takara Bio. Hygromycin B and protease inhibitors were from Roche. Odyssey Blocking Buffer was from LI-COR. Gene amplification primers were from Integrated DNA Technologies. Phusion polymerase, restriction enzymes, T4 DNA ligase and the Gibson Assembly Cloning Kit were from New England Biolabs.

### Antibodies

Primary monoclonal antibodies used and their sources (indicated in parentheses) were: mouse anti-TYRP1 (TA99/Mel-5, American Type Culture Collection; Rockville, MD); mouse anti-γ-Tubulin (GTU-88, Sigma); mouse anti-γ-adaptin (BD 610385); mouse anti-GFP (clones 7.1 and 13.1, Roche); rabbit anti-AP3M1 (ab201227; Abcam); rat anti–mouse LAMP2 (GL2A7, Abcam); rat anti-mouse TfR (CD71, BD 553264); and rat anti-HA (3F10, Roche 11867423001). Primary polyclonal antibodies used and their sources were: rabbit anti-STX13 (a kind gift of Rytis Prekeris, Univ. of Colorado, Denver, CO, USA)(Prekeris et al., 1998); rabbit anti-pallidin (a kind gift of Juan Bonifacino, National Institute of Child Health and Human Development, Bethesda, MD, USA)(Moriyama and Bonifacino, 2002); rabbit anti-PI4KIIIβ (13247-1-AP, Proteintech); rabbit anti-PI4KIIα and anti-PI4KIIβ (kind gifts of Pietro De Camilli, Yale Univ., New Haven, CT, USA) (Guo et al., 2003); and rabbit anti-GFP (PABG1, Chromotek); Species- and/or mouse isotype–specific secondary antibodies from donkey or goat conjugated to Alexa Fluor 488, Alexa Fluor 594, or Alexa Fluor 640 used for IFM or to Alexa Fluor 680 or Alexa Fluor 790 for immunoblots were obtained from Jackson ImmunoResearch Laboratories.

### DNA constructs

All oligonucleotides used in constructing shRNA and expression plasmids are documented in Supplementary Table 1. The pLKO.1-puromycin derived lentiviral vectors for small hairpin RNAs (shRNAs) against mouse PI4KIIα (shPI4KIIα-1, shPI4KIIα-2), PI4KIIβ (shPI4KIIβ-1, shPI4KIIβ-2), pallidin (shPallidin-1, shPallidin-2), PI4KIIIβ (shPI4KIIIβ-1, shPI4KIIIβ-2) and non-target control (shNC) were obtained from the High-throughput Screening Core of the University of Pennsylvania (Supplementary Table1). The plasmid-based expression vectors pEGFP-N1-PI4KIIα (Balla et al., 2002), pEGFP-N1-PI4KIIβ (Balla et al., 2002), pEGFP-C1-P4C-SidC (Dolinsky et al., 2014; Weber et al., 2018; Weber et al., 2014), pEGFP-C1-VAMP7 (Dennis et al., 2016), pmCherry-STX13 (Dennis et al., 2016; Dennis et al., 2015), pEGFP-C3-2xFYVE (Gillooly et al., 2000) and pCR-OCA2-WT-HA-UTR2 (Sitaram et al., 2009) and the retroviral expression vector pBMN-mCherry-STX13-IRES-Hygro (X/N) (Bowman et al., 2021) have been previously described. Lentiviral packaging plasmid psPAX2 and pMD2.G encoding the vesicular stomatitis virus glycoprotein were gifts from Didier Trono (Univ. of Geneva) and purchased from Addgene (http://n2t.net/addgene:12260 and http://n2t.net/addgene:12259, respectively). Plasmid pBMN-GFP-P4C-SidC was generated by amplifying GFP-P4C-SidC from pEGFP-C1-P4C-SidC using primers 1 and 2, subcloning the amplified fragment into the NotI restriction site of pBMN-IRES-hygro(X/N) (Setty et al., 2007), and determining the proper insert orientation by gene amplification. The shRNA-resistant (shPI4KIIα-1^R^) forms of PI4KIIα-WT-GFP, PI4KIIα-D308A-GFP and PI4KIIα-L61, 62A-GFP were generated by two-step gene amplification using the previously described MiGR-PI4KIIα-WT-GFP, MiGR-PI4KIIα-D308A-GFP and MiGR-PI4KIIα-L61, 62A-GFP, respectively, as templates (López-Haber et al., 2020). In the first step, fragments containing PI4KIIα cDNA 1-477 and PI4KIIα cDNA 459-1437 fused to GFP were separately amplified using primers 3 and 4 and primers 5 and 6, respectively. The two PCR products contain a 19-bp region of overlap containing the shPI4KIIα-1^R^ sequence CAAaAATGAgGAaCCCTAT (mutagenized sites in small caps). The second step amplification concatenated the two PCR products using primers 3 and 6 to generate the full-length products, which were then subcloned into the NotI restriction site of pBMN-IRES-hygro(X/N) to generate pBMN-PI4KIIα-WT-GFP, pBMN-PI4KIIα-D308A-GFP and pBMN-PI4KIIα-L61, 62A-GFP.

Similarly, the shRNA-resistant (shPI4KIIβ-2^R^) forms of PI4KIIβ-WT-GFP and PI4KIIβ-D304A-GFP were generated by two-step amplification using pEGFP-N1-PI4KIIβ-WT and pEGFP-N1-PI4KIIβ-D304A (Balla et al., 2002) as templates, respectively. In the first step, fragments containing PI4KIIβ cDNA bp 1-1080 and PI4KIIβ bp cDNA 1063-1443 fused to GFP were separately amplified using primers 7 and 8 and primers 9 and 6, respectively. The two PCR products contain an 18-bp region of overlap containing the shPI4KIIβ-2^R^ sequence GAcGAgTGGAGAGCtTAT (mutagenized sites in small caps). The second step amplification concatenated the two PCR products using primers 7 and 6 to generate the full-length products, which were then subcloned into the NotI restriction site of pBMN-IRES-hygro(X/N) to generate pBMN-PI4KIIβ-WT-GFP and pBMN-PI4KIIβ-D308A-GFP. In all cases, the proper orientation of inserts within the pBMN-IRES-hygro vector was determined by gene amplification. The shPI4KIIβ-2^R^ form of pBMN-PI4KIIβ-L29, 30A-GFP was made by Gibson assembly using pBMN-PI4KIIβ-WT-GFP as template. Fragments containing PI4KIIβ cDNA bp 1-1017 and cDNA bp 987-1443 fused to GFP were amplified using primers 10 and 11, and primers 12 and 13, respectively. The two PCR products contain a 28-bp region of overlap GAGCC**GgcGgc**ACCGCGGATCGCCTGGG in which the codons for the two leucines of the AP-1 binding site are mutagenized to alanine codons (**alanine codons** in bold).

### Cell culture and cell line generation

Immortalized melanocyte cell lines melan-Ink4a-1 (referred to here as melan-Ink4a or WT) derived from C57BL/6J-*Ink4a-Arf^-/-^ (Cdk2a*-null) mice (Sviderskaya et al., 2002) and melan-pa1 (referred to here as melan-pa) derived from Pldn/BLOC-1-deficient C57BL/6J-*Pldn^pa/pa^* pallid mice (Setty et al., 2007) have been described. All melanocyte cell lines were cultured at 37°C and 10% CO_2_ in RPMI 1640 supplemented with 10% FBS (Atlanta Biologicals), 2 mM L-glutamine and 200 nM 12-*O*-tetradecanoylphorbol-13-acetate. Plat-E cells (Morita et al., 2000) were cultured at 37°C and 5% CO_2_ in DMEM supplemented with 10% FBS (Hyclone), 2 mM L-glutamine, 1 mM sodium pyruvate with 1 μg/mL puromycin and 10 μg/mL blasticidin to maintain selection for the murine leukemia virus Gag and Env genes. HEK293T cells were cultured at 37°C and 5% CO_2_ in DMEM supplemented with 10% FBS (Hyclone), 2 mM L-glutamine. All cell lines were tested to be negative for mycoplasma every 2–3 months using the MycoAlert Mycoplasma Detection Kit (Lonza).

Stable melan-Ink4a lines expressing mCherry-tagged STX13 or GFP-tagged P4C-SidC, PI4KIIα or PI4KIIβ were generated by transducing melan-Ink4a cells with recombinant retroviruses encoding the indicated genes. Retroviruses were produced by Plat-E cells that were transiently transfected with indicated retroviral DNA constructs in the pBMN-IRES-hygro vector. Briefly, Plat-E cells were plated at 1.5×10^6^ cells per 35-mm well and transfected the next day with 3.5 μg plasmid DNA using Lipofectamine 2000 (Thermo Fisher) in DMEM-containing medium according to manufacturer’s instructions. The medium was replaced with RPMI 1640/ 10% FBS after 4-6 h, and retrovirus-containing supernatants were collected 48 h later, filtered, and added to melan-Ink4a cells seeded the previous day (3-4×10^5^ cells per 35-mm well). The retrovirus-containing supernatants was replaced with fresh medium containing 400-500 μg/ml hygromycin B, and polyclonal drug-resistant lines were selected over 14 d. Stable transfectant lines were occasionally treated with 200 µg/ml hygromycin B for 2–3 d to maintain selective pressure for the transgene.

Lentiviruses encoding shRNAs were harvested from transfected HEK293T cells that were co-transfected with packaging plasmid psPAX2, envelope plasmid pMD2.G, and pLKO.1 lentiviral plasmid containing the indicated shRNA insert and puromycin-resistance gene. HEK293T cells plated at 1.5×10^6^ cells per 35-mm well the day before were transfected with 3.5 μg pLKO.1-shRNA, 2.625 μg psPAX2, and 0.875 μg pMD2.G using Lipofectamine 2000 (Thermo Fisher) according to the manufacturer’s instructions. Medium replacement and virus harvesting were as described above for retroviral production. The lentivirus-containing supernatants were diluted 1:1 with fresh medium and added to melanocytes seeded the previous day (3-4×10^5^ cells per 35-mm well). Transduced cells were selected using 2 μg/mL puromycin for 7-14 d or as indicated otherwise.

For transient transfection of melanocytes, cells were plated at 2.5-3×10^6^ cells per 35-mm well on Matrigel (BD)-coated coverslips (for OCA2-HA, GFP-VAMP7) or 35-mm glass-bottomed dishes (Matsunami D35-14-1.5-U; for mCherry-STX13, GFP-P4C, PI4KIIα-GFP, PI4KIIβ-GFP, GFP-2XFYVE) and then transfected the next day with 1 μg (GFP-P4C, GFP-2XFYVE) or 2 μg (OCA2-HA, GFP-VAMP7, mCherry-STX13, PI4KIIα-GFP, PI4KIIβ-GFP) of expression plasmid using Lipofectamine 3000 according to the manufacturer’s instructions. Cells were imaged 24 h (GFP-VAMP7, GFP-P4C, PI4KIIα-GFP, PI4KIIβ-GFP, GFP-2XFYVE) or 48 h (OCA2-HA, mCherry-STX13) after transfection.

### Melanin content quantification

Quantification of melanin content by spectroscopy was performed essentially as described (Delevoye et al., 2009). Briefly, 1.2×10^6^ cells were seeded in 6-cm dishes in triplicate. The next day, cells were rinsed twice with PBS, detached by incubation in PBS, 10 mM EDTA for 15-20, and collected by centrifugation at 500 x *g* for 5 min. Cells were resuspended in cold 50 mM Tris-HCl, pH 7.4, 2 mM EDTA, 150 mM NaCl, 1 mM DTT, and 1x protease inhibitor cocktail (Roche), and then sonicated on ice using the Sonic Dismembrator Model 100 (Fisher Scientific). Sonicates were fractionated into supernatant and pellet fractions by centrifugation at 20,000 x *g* for 15 min at 4°C. Supernatants were subjected to determination of protein concentration by BCA protein determination kit (Thermo Fisher). Melanin-containing pellets were rinsed in ethanol/diethyl ether (1:1) and dissolved in 2 M NaOH/ 20% dimethyl sulfoxide at 60°C for 1 hour. The melanin content of individual samples was measured by spectrophotometry as absorbance at 492 nm, and normalized to protein concentration in each sample. Triplicate samples were analyzed for each experimental condition in each experiment.

### Immunoblotting

Melanocytes were cultured in 10-cm dishes, detached by incubation in PBS, 10 mM EDTA for 15-20 min, and collected by centrifugation at 500 x *g* for 5 min. Cell pellets were resuspended in cold 50 mM Tris-HCl, pH 7.4, 2 mM EDTA, 150 mM NaCl, 1 mM DTT, and 1x protease inhibitor cocktail (Roche), and lysed by sonication on ice using the Sonic Dismembrator Model 100 (Fisher Scientific). Concentrated Laemmli sample buffer with 2-mercaptoethanol was added to cell lysates and heated at 95-100°C for 10 min. Samples were then fractionated by 12% sodium dodecyl sulfate (SDS)/polyacrylamide gel electrophoresis (PAGE) on polyacrylamide gels and transferred to polyvinylidene difluoride membranes (Immobilon-FL, Millipore). Membranes were blocked with Odyssey Blocking Buffer at room temperature (RT) for 1 h, incubated with primary antibody diluted in TBS/ 0.2% tween-20 (TBST) overnight at 4°C, washed 3 times 10 min with TBST, and then incubated with secondary antibody conjugated with either Alexa Fluor 680 or Alexa Fluor 790, diluted in TBST, at RT for 1 h. After washing 3 times 10 min with TBST, blots were analyzed using a LI-COR Odyssey CLx imaging system, and images were further processed and quantified using Fiji-ImageJ (National Institutes of Health).

### Subcellular fractionation

Subcellular fractionation was performed essentially as described (Di Pietro et al., 2004; Di Pietro et al., 2006) with minor modifications. All steps were performed at 4°C. To prepare cytosolic and membrane fractions, melanocytes were homogenized at 10^7^ cells/ml in hypotonic buffer [10 mM HEPES, pH 7.4, 1 mM MgCl_2_, 1 mM EGTA, 250 mM sucrose, 2 mM DTT, 1x protease inhibitor cocktail (Roche), 1x phosphatase inhibitor (Roche)] by passing cells through a pre-chilled 28G needle for 25-30 cycles until 70-80% cells were disrupted as evaluated by trypan blue exclusion. Undisrupted cells in the crude homogenate were pelleted by centrifugation at 500 x *g* for 1 min, and the supernatant was then centrifuged at 5000 x *g* for 10 min to remove nuclei and mitochondria. The resulting post-nuclear supernatant was fractionated into membrane pellet and cytosolic supernatant fractions by centrifugation at 100,000 x *g* for 60 min. Membrane pellets were gently washed once in hypotonic buffer to minimize contamination by cytosolic proteins and then solubilized in hypotonic buffer containing 0.2% SDS to the same volume as the cytosol fraction. 6X Laemmli sample buffer with 2-mercaptoethanol was added to solubilized membrane and cytosolic fractions (to a final 1X concentration), and samples were heated at 95-100°C for 10 min. Identical volumes of membrane and cytosolic fractions were fractionated by SDS-PAGE on 12% polyacrylamide gels and analyzed by immunoblotting.

### Immunofluorescence microscopy and live-cell imaging

Immunofluorescence microscopy was performed essentially as described (Bowman et al., 2021; Dennis et al., 2016). Briefly, melanocytes were seeded on matrigel-coated coverslips 24-48 h before fixation. Cells were fixed with 4% formaldehyde (VWR Scientific) in PBS (RT, 15 min), washed 5 times with PBS, incubated with primary antibody (RT, 1 h) diluted in blocking buffer (0.01% BSA and 0.02% Saponin in PBS), washed 3 times 5 min with PBS, incubated with secondary antibodies (RT, 30 min) diluted in blocking buffer, washed again, and mounted onto slides using Prolong Gold (Thermo Fisher). Fixed cell images were acquired using a Leica DMI8 inverted microscope (Leica Biosystems) equipped with a CSU-WI spinning-disk confocal scanner unit (Yokogawa), a 100X total internal reflection fluorescence objective lens (Leica 1.47 NA) or a 63X objective lens (Leica, 1.40 NA) a Hamamatsu Photonics ORCA-Flash 4.0 sCMOS digital camera, and VisiView software (Visitron System). Images were further processed and analyzed using Fiji-ImageJ.

Live-cell imaging was done essentially as described (Bowman et al., 2021). Briefly, melanocytes were seeded onto matrigel-coated 35-mm glass-bottomed dishes, incubated in Leibovitz’s CO_2_-independent medium without phenol red (Invitrogen) supplemented with 10% FBS in an environmental chamber set to 37°C, and imaged using the same microscopic system. To image tubule dynamics, z-series images spanning two to five layers separated by 0.2-μm or 0.25-μm were captured every 0.6–1 s for a minimum of 30 s and a maximum of 90 s. To observe cellular PtdIns4P distribution, sixteen 0.2-μm layers of z-series were acquired for further analysis. Dual-view live-cell imaging was performed using the Hamamatsu W-VIEW GEMINI Image splitting optics, and one to three z-series layers separated by 0.2-μm were captured every 0.4-0.5 s for a minimum of 20 s and a maximum of 80 s.

### Quantitative image Analysis

#### Colocalization analyses

The area of overlap between fluorescently labeled proteins and pigmented melanosomes or between two fluorescently labeled proteins was quantified using Fiji-ImageJ (Schneider et al., 2012) on fixed cell images essentially as previously described (Bowman et al., 2021; Dennis et al., 2015). Briefly, single z-plane images were cropped around single cells, and the nucleus and densely labeled perinuclear region were removed. The local background of fluorescent images was subtracted using the “rolling ball” algorithm with “sliding paraboloid” selected; the rolling ball radius was chosen based on fluorescence intensity, such that a smaller radius was used for higher fluorescence intensity values (e.g. 2 pixels for most TYRP1, STX13, LAMP2, OCA2-HA, and PI4KIIα images, and 5 pixels for TfR, PI4KIIβ, and PI4KIIα-GFP) and a higher radius for lower intensity values (e.g. 10 pixels for GFP-VAMP7 and PI4KIIβ-GFP). Pigment granules from the bright field channel were processed using either the “Invert” function first and then “Threshold” function by ImageJ or, for Figure 1C, the plugin “Spot Detector” by Icy software (https://icy.bioimageanalysis.org/). After background subtraction, images were converted to binary images using either the “Threshold” function (TYRP1, STX13, LAMP2 and OCA2-HA) or the Bernson algorithm of the “Auto Local Threshold” function with a 15-pixel radius (GFP-VAMP7, PI4KIIα, PI4KIIβ, PI4KIIα-GFP, PI4KIIβ-GFP) or both (TfR).

Structures of less than 5 pixels in area were considered background and filtered out using the “Analyze Particles” function. To generate an overlap image, two binary images were multiplied using the “Image Calculator” function, and the areas of the individual and overlap binary images were calculated using the “Analyze Particles” function. Percentage of overlap was calculated either as the percentage of the area of overlap relative to the area of one of the individual labels or by dividing the number of TYPR1-overlapping pigment granules by the total number of pigment granules (**Fig. 1C**).

#### Tubule number and percentage of STX13-positive tubules that contain PtdIns4P

Quantification of tubules containing PtdIns4P, as labeled by the GFP-P4C-SidC (P4C) probe, was done using ImageJ on video streams of melan-Ink4a mouse melanocytes that stably express P4C and transiently express mCherry-STX13 (STX13). A maximum-intensity projection was generated from three consecutive z-series layers separated by 0.2-μm from a single frame of a video stream, and two cytoplasmic 100-μm^2^ square regions were randomly selected for each single cell analysis. Background subtraction and segmentation of both P4C and STX13 were processed using the SQUASSH algorithm (Rizk et al., 2014) with the following parameters: rolling ball window size of 20 pixels in the Background subtractor; 0.04 for minimum object intensity channel; “Subpixel segmentation” selected; and 0.4 for standard deviation of xy and z in the PSF model. Binary images were generated using the “SQUASSH” algorithm, and structures in which binary P4C- and STX13-containing structures overlapped were visualized using the “Image Calculator” “multiply” function. Circular structures in binary images were removed and P4C-positive and/or P4C-STX13-positive tubular structures were quantified using the Analyze particles function with size of 0.0625 μm^2^-infinity and circularity of 0-0.7. The percentage of STX13-positive tubules that overlap P4C was calculated by dividing the number of P4C-STX13-positive tubules by the number of STX13-positive tubules. In the graphs shown in **Fig. 8B-D**, a single dot indicates the average of two 100-μm^2^ regions within a single cell.

#### Number, percentage, and length of STX13-containing tubules

Quantification of endosomal tubules labeled by mCherry-STX13 was performed using Fiji-ImageJ on melan-Ink4a mouse melanocytes that stably express mCherry-STX13. Single plane images with largely in-focus STX13-positive structures were selected from image sequences, and ten cytoplasmic 25-μm^2^ square regions were randomly chosen and analyzed individually. Signal background was subtracted using the the Rolling Ball algorithm with a 10-pixel radius and Sliding Paraboloid selected. After background subtraction, images were processed by Smooth to reduce noise and then rendered binary using the Phansalkar algorithm of the “Auto Local Threshold” function with a 15-pixel radius, followed by the Skeletonize function. The Analyze Skeleton function was used to calculate and document the length and number of tubules, represented as Branches. Only branches longer than 0.25 μm were considered as tubules. Tubule length was determined by averaging all branches longer than 0.25 μm within each 25-μm^2^ region of a single cell. In the graphs shown in **Fig. 9D-G**, a single dot represents the average of ten 25-μm^2^ regions within a single cell.

#### Number of GFP-2XFYVE-containing puncta per cell

Quantification was done using ImageJ from video streams of melan-Ink4a mouse melanocytes expressing shNC, shPI4KIIα, or shPI4KIIβ. A maximum-intensity projection of selected frames was generated from three consecutive z-series layers separated by 0.25-μm z-step, and regions covering entire cells were analyzed. Signal background was subtracted using the Rolling Ball algorithm with a 5-pixel radius with Sliding Paraboloid selected. After background subtraction, images were rendered binary using the “Contrast” algorithm of the “Auto Local Threshold” function with a 15-pixel radius, followed by the “Fill holes” and “Watershed” functions. Using the “Analyze Particles” function, structures of less than 5 pixels were considered background and filtered out, and the number of puncta in each cell was calculated.

#### Graphical presentation of data and statistical analyses

At least three independent replicates were performed in each experiment and sample sizes are as indicated in each figure legend. In all graphs, results were presented as mean ± SD. Most data are presented as “super-plots” (Lord et al., 2020) in which values from individual cells within each experiment are indicated by a single dot, colors represent independent experiments, triangles represent means of each experiment, and bars represent the overall mean and SD across all experimental repeats. Statistical analyses were performed in GraphPad Prism using ordinary one-way ANOVA, Welsh’s ANOVA, or Kruskal-Wallis test with post-hoc correction as specified and determined using GraphPad Prism by assessing each data set for normality and homoscedasticity; Welsh’s ANOVA was used when the data set did not display homoscedasticity, and a Kruskal-Wallis test was used if the data set did not display a normal distribution. Significance between control or experimental samples are indicated; only *p* < 0.05 was considered as statistically significant.

## Supplemental Data

Supplemental Figure S1 shows target protein levels after 7-9 d shRNA treatment and the effect of PI4KIIIβ depletion on TYRP1 localization to melanosomes. Supplemental Figure S2 shows representative images of the pigmentation of cells treated with shRNAs to PI4KIIα, PI4KIIβ, or Pallidin after 14 d, and documents TYRP1 localization to melanosomes following long-term (30 d) PI4KIIα or PI4KIIβ shRNA treatment. Supplemental Figure S3 shows that 7-9 d depletion of either PI4KIIα or PI4KIIβ by shRNA in WT melanocytes impairs localization of GFP-VAMP7 to melanosomes. Supplemental Figure S4 shows validation of TfR as an early endosomal marker in WT and BLOC-1^-/-^ melanocytes. Supplemental Figure S5 shows that the distribution of PtdIns3P is not affected by 7-9 d treatment with PI4KIIα or PI4KIIβ shRNA.

## Abbreviations

AP-1: adaptor protein-1

AP-3: adaptor protein-3

BLOC: Biogenesis of Lysosome-related Organelles Complex

HPS: Hermansky-Pudlak syndrome(s)

IFM: immunofluorescence microscopy

LROs: lysosome-related organelles

NC: non-target control

OCA2: oculocutaneous albinism type 2 protein

P4C: *Legionella pneumophila* P4C-SidC

PI4K: phosphatidylinositol-4-kinase

PtdIns: phosphatidylinositol

PtdIns4P: phosphatidylinositol-4-phosphate

STX13: syntaxin-12/13

VAMP7: Vesicle Associated Membrane Protein 7

TfR: transferrin receptor

TYRP1: tyrosinase-related protein-1

WT: wild-type

## ACKNOWLEDGMENTS

We thank Tamas Balla, Sergio Grinstein, and Christopher Burd for PI4KIIα and PI4KIIβ cDNAs, the P4C probe, and helpful discussions, Pietro De Camilli, Juan Bonifacino, and Rytis Prekeris for critical antibodies, Shanna Bowman and Linh Le for advice on video microscopy and image quantification, Dawn Harper for laboratory assistance, and Leslie King for helpful comments. We are grateful for funding from National Institutes of Health grants R01-EY015625 from the National Eye Institute (to MSM and GR), Institut National de la Santé et de la Recherche Médicale (to CD), Centre National de la Recherche Scientifique (to GR), Institut Curie (to GR and CD), and Genespoir (to CD).

## AUTHOR CONTRIBUTIONS

All authors edited the manuscript. In addition:

<u>Yueyao Zhu</u> conceptualized the project and experimental design, performed most of the experiments, analyzed most of the data, assembled the figures, wrote the original draft of the paper, and contributed to final editing of the manuscript.

<u>Shuixing Li</u> contributed to experimental design and performed some experiments.

<u>Alexa Jaume</u> contributed to experimental design, performed some experiments, and analyzed data.

<u>Riddhi Atul Jani, Cédric Delevoye, and Graça Raposo</u> contributed to conceptualization of the project, experimental design, and data interpretation.

<u>Michael S. Marks</u> conceptualized, supervised, and administered the project, coordinated among all contributors, provided resources, analyzed data, edited figures, wrote part of the original manuscript draft, and coordinated editing by all co-authors.

## Notes

### Competing Interest Statement

The authors have declared no competing interest.

### Summary of Updates

All figures and supplemental figures have been revised; new control data have been added; new quantification of previous data has been added; statistics have been updated; text has been revised and edited for clarity.

