## Supplemental for "Two type II phosphatidylinositol 4-kinases function sequentially in tubule-mediated cargo delivery from early endosomes to melanosomes"

**Zhu et al.**

**SUPPLEMENTAL DATA**

Supplemental Figures S1-S5

Supplemental Table S1

**SUPPLEMENTAL FIGURES**

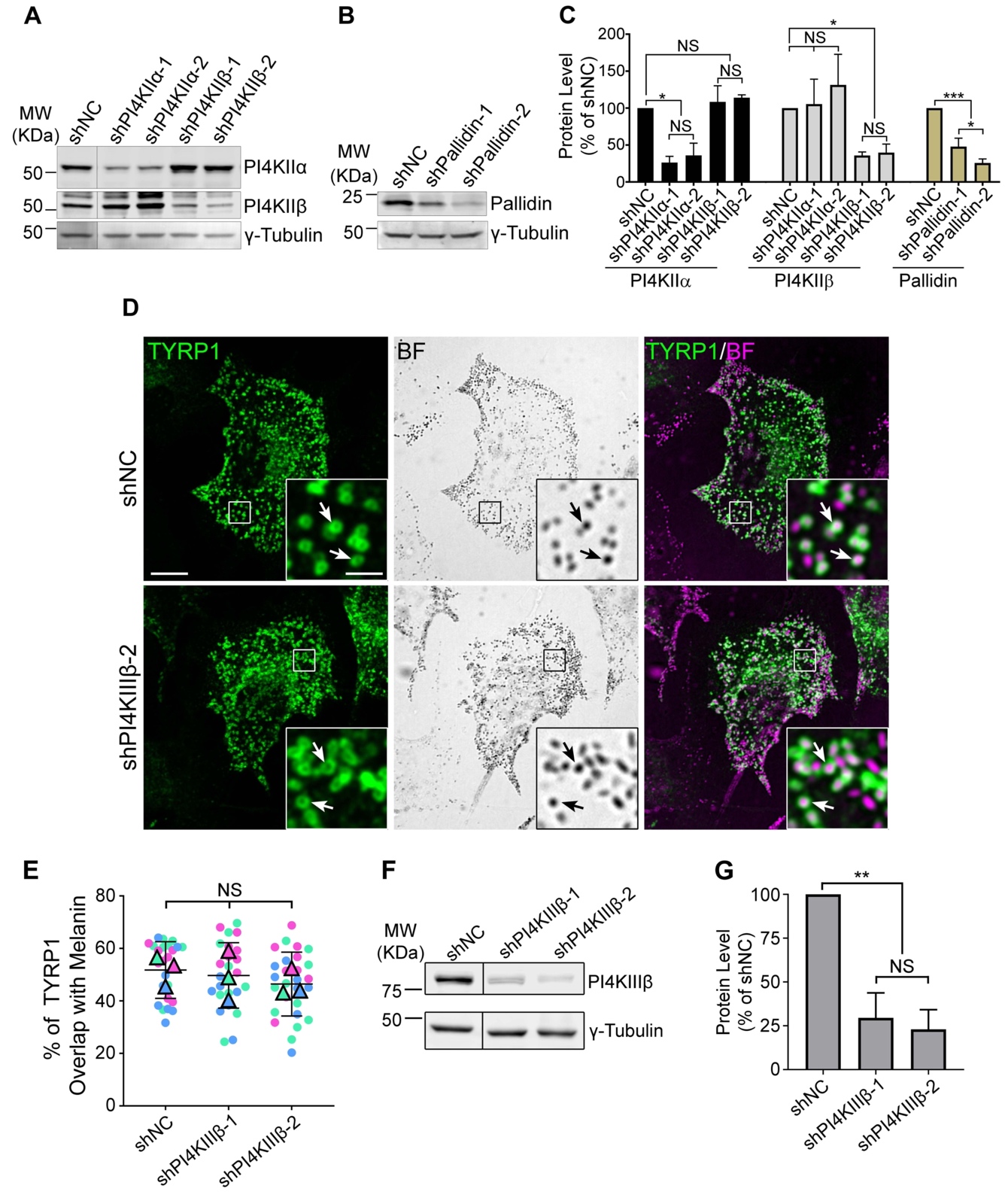

**Figure S1. Target protein levels after 7-9d shRNA treatment and TYRP1 localization to melanosomes in cells depleted of PI4KIIIβ.** WT melan-Ink4a melanocytes were transduced with lentiviruses expressing the indica ted shRNAs to PI4KIIα, PI4KIIβ, PI4KIIIβ, or the Pallidin subunit of BLOC-1, or control non-coding shRNA (shNC) and selected for 8 d. **(*A***, ***B***, ***F*)** Whole-cell lysates from cells treated with the indicated shRNAs were analyzed by SDS/PAGE and immunoblotting for PI4KIIα, PI4KIIβ, PI4KIIIβ, or Pallidin and for γ-Tubulin as a loading control. Positions of the 25 and 50 kDa molecular weight (MW) markers (A, B) or the 75 and 50 kDa MW markers (F) are indicated. Note that PI4KIIβ migrates as 2-3 distinct bands, all of which are depleted upon treatment with specific shRNA. **(*C*** and ***G*)** Quantification of band intensities (mean ± SD) for the indicated components normalized to γ-Tubulin from three independent experiments each, and analysis by one-way ANOVA with repeated measures. NS, not significant; *, p<0.05; **, p<0.01; ***, p<0.005.  **(*D*)** Cells at 9 days post-infection with either of two distinct shRNAs to PI4KIIIβ were analyzed for endogenous TYRP1 by IFM (green, left and right panels) and for pigment granules by bright field microscopy (BF; middle panels and pseudocolored magenta in the merged images on the right). White arrows, examples of TYRP1 surrounding pigment granules; boxed regions are magnified 5-fold in the insets. Scale bars: main panels, 10 μm; insets, 2 μm. **(*E*)** Quantification of TYRP1 overlap with melanosomes. Data from three independent experiments each were analyzed by ordinary one-way ANOVA; NS, not significant.

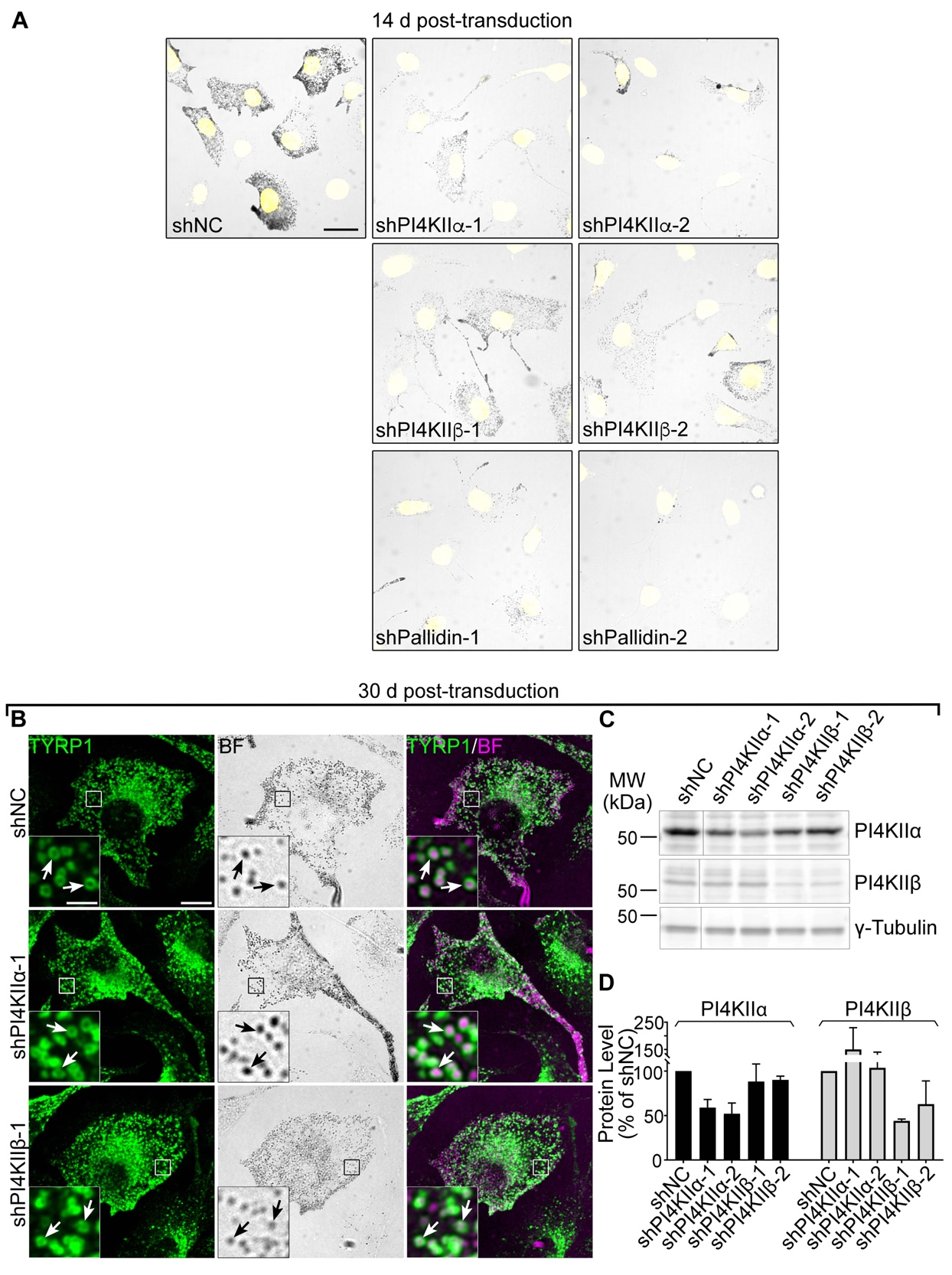

**Figure S2. Pigmentation of cells after 14 d treatment with shRNAs to PI4KIIα, PI4KIIβ, or Pallidin and TYRP1 localization to melanosomes following 30 d PI4KIIα or PI4KIIβ shRNA treatment.**  **(*A*)** WT melan-Ink4a melanocytes were transduced with lentiviruses expressing the indicated shRNAs to PI4KIIα, PI4KIIβ, PI4KIIIβ, or the Pallidin subunit of BLOC-1, or control non-coding shRNA (shNC) and selected for 14 d as in **Fig. 1F**. Fixed cells were labeled with Hoechst 33342 dye and analyzed by fluorescence (for Hoechst; pseudocolored yellow) and bright field microscopy (for pigment). Shown are representative images documenting the pigmentation status of the cells. Scale bars, 20 μm. **(*B***-***D*)** WT melan-Ink4a melanocytes were transduced with the indicated shRNAs to PI4KIIα or PI4KIIβ and analyzed 30 d post-infection. (B) Cells were analyzed for endogenous TYRP1 by IFM (green, left and right panels) and for pigment granules by bright field microscopy (BF; middle panels and pseudocolored magenta in the merged images on the right). White arrows, examples of TYRP1 surrounding pigment granules; Boxed regions are magnified 5-fold in the insets. Scale bars: main panels, 10 μm; insets, 2 μm. (C) Whole-cell lysates from cells treated with the indicated shRNAs were analyzed by SDS/PAGE and immunoblotting for PI4KIIα, PI4KIIβ, or Pallidin and for γ-Tubulin as a loading control. Positions of the 25 and 50 kDa molecular weight (MW) markers are indicated. Note that PI4KIIβ migrates as 2-3 distinct bands, all of which are depleted upon treatment with specific shRNA. (D) Quantification of band intensities (mean ± SD) for the indicated components normalized to γ-Tubulin from two independent experiments.

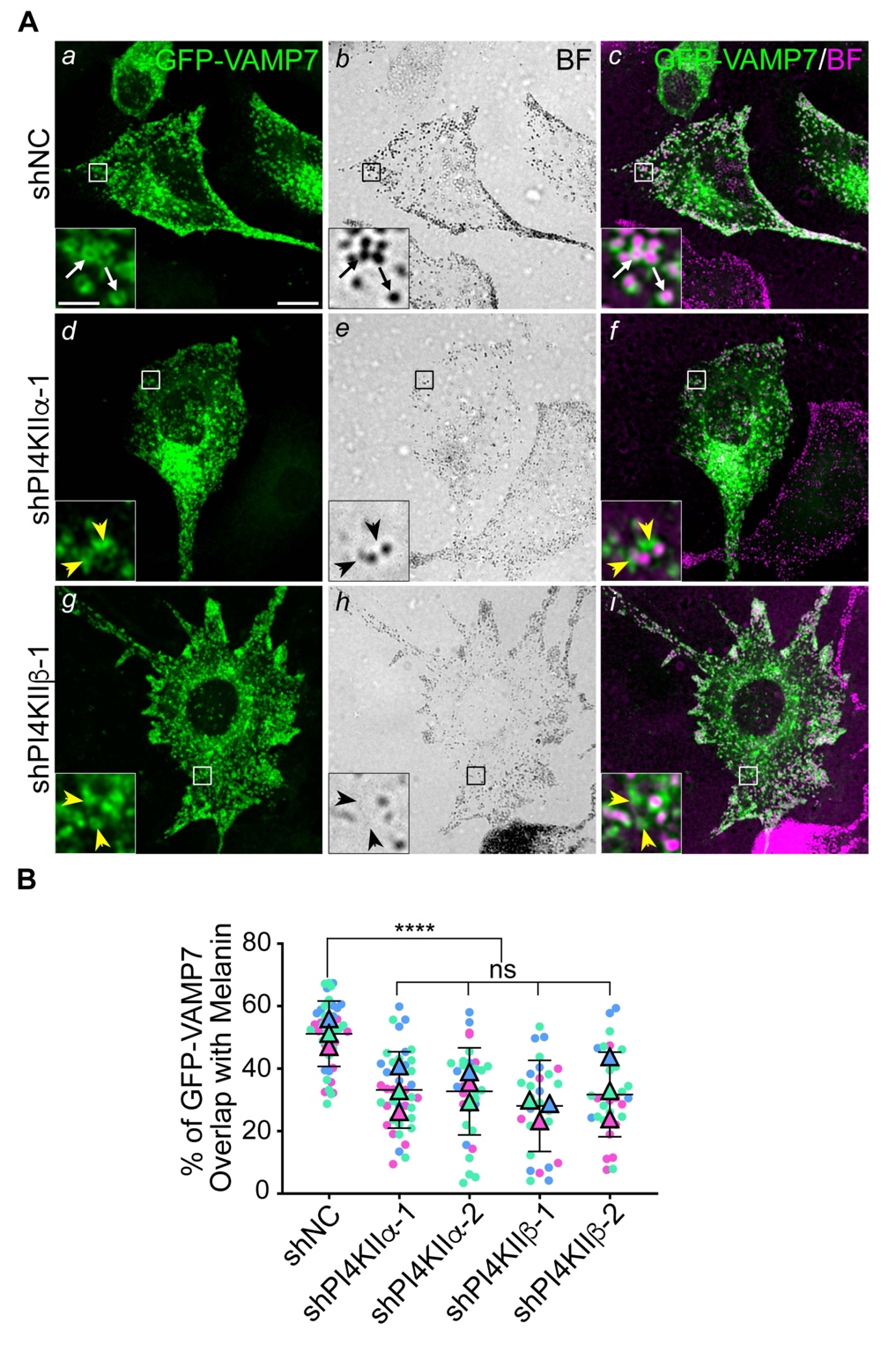

**Figure S3. Depletion of either PI4KIIα or PI4KIIβ by shRNA in WT melanocytes impairs localization of GFP-VAMP7 to melanosomes.** WT melan-Ink4a melanocytes were transduced with lentiviruses expressing the indicated shRNAs to PI4KIIα, PI4KIIβ, or control non-coding shRNA (shNC) and selected for 7 to 9 d. **(*A*)** After shRNA transduction, cells were transiently transfected with GFP-VAMP7 and analyzed for GFP-VAMP7 by IFM (green, left and right panels) and for pigment granules by bright field microscopy (BF; middle panels and pseudocolored magenta in the merged images on the right). White arrows, examples of GFP-VAMP7 surrounding pigment granules; yellow arrowheads, examples of mislocalized GFP-VAMP7 not associated with pigment granules. Boxed regions are magnified 5-fold in the insets. Scale bars: main panels, 10 μm; insets, 2 μm. **(*B*)** Quantification of the overlap of GFP-VAMP7 with pigment granules in cells treated with the indicated shRNAs. Data from three independent experiments were analyzed by ordinary one-way ANOVA. ****, p<0.0001; ns, not significant.

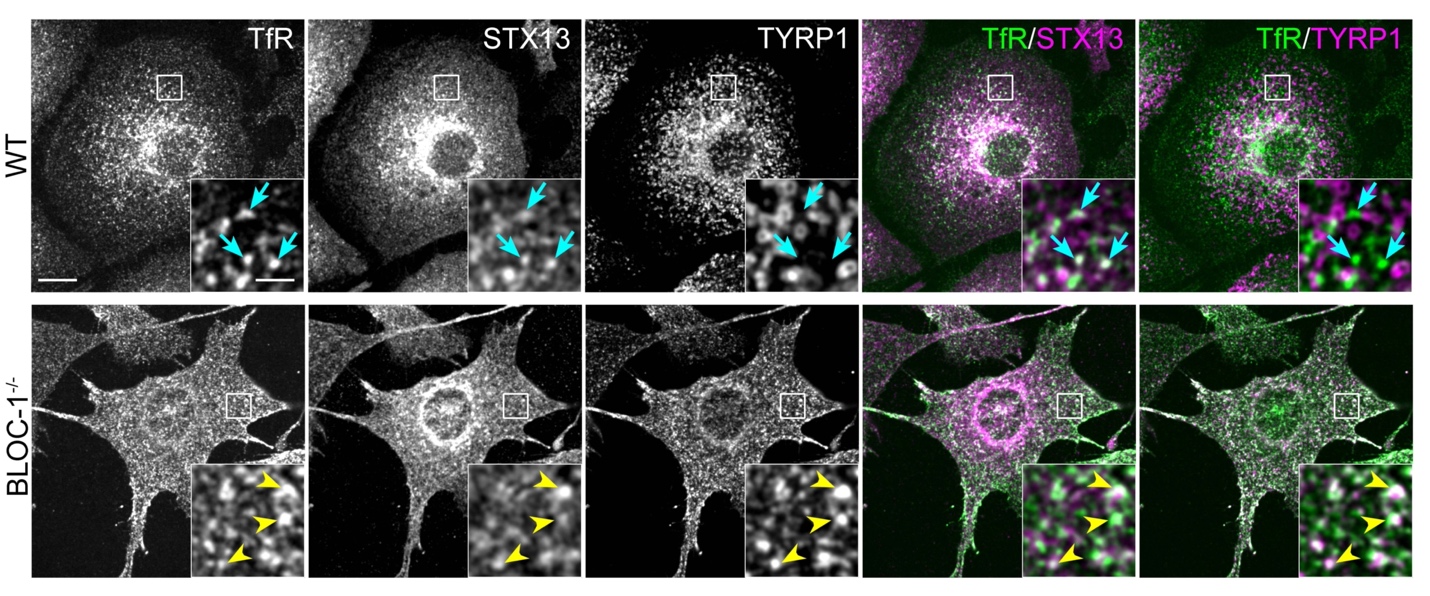

**Figure S4. Validation of TfR as an early endosomal marker.** WT melanocytes (upper panels) and BLOC-1^-/-^ melanocytes from pallid mice (lower panels) were fixed and analyzed by triple label IFM for endogenous TfR (white and green in single and merged panels, respectively) relative to STX13 and TYRP1 (white and magenta in single and merged panels, respectively). Cyan arrows, examples of TfR overlap with STX13 but not TYRP1; yellow arrowheads, examples of TfR overlap with both STX13 and TYRP1. Boxed regions are magnified 5-fold in the insets. Scale bars: main panels, 10 μm; insets, 2 μm.

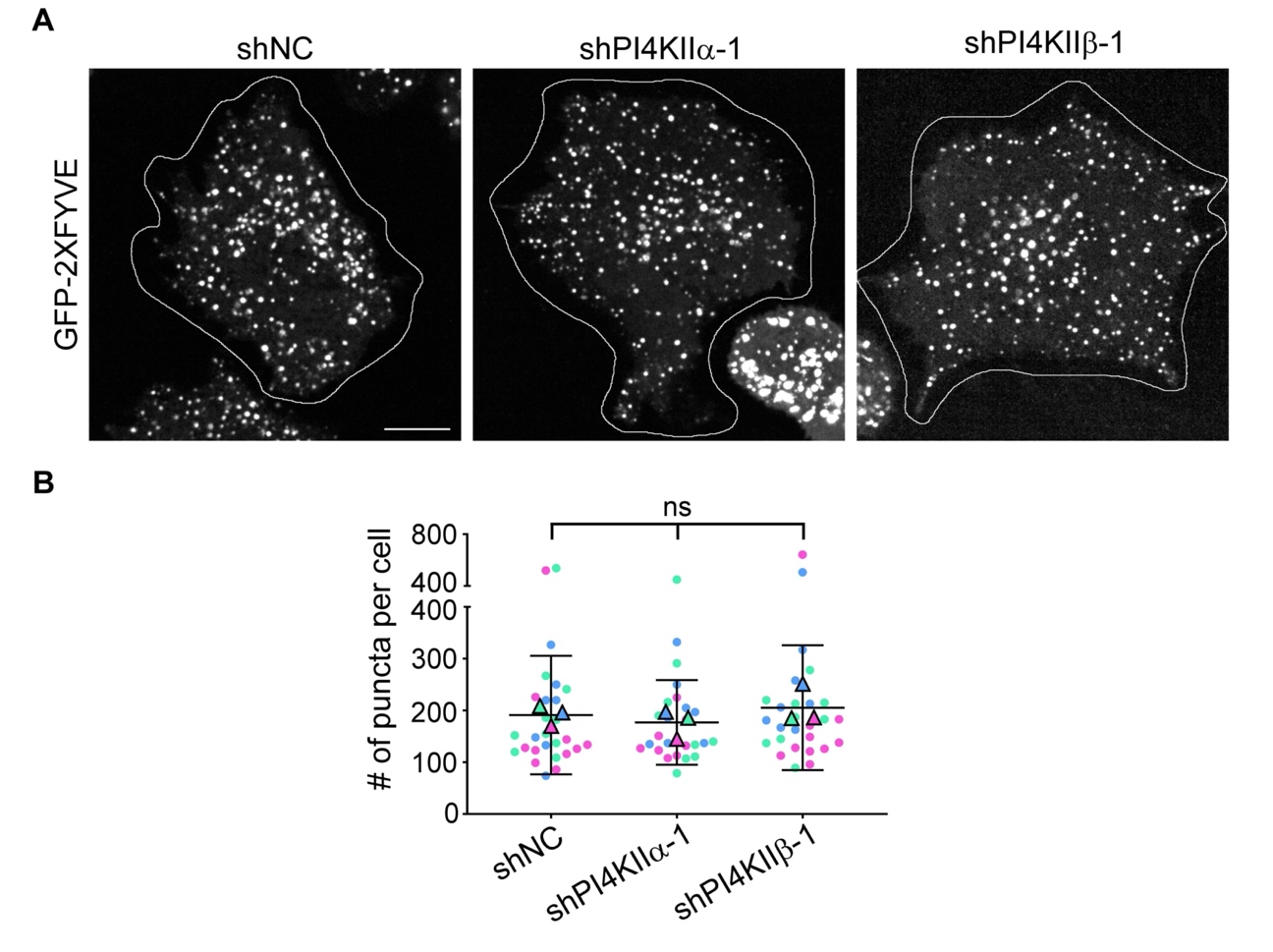

**Figure S5. The distribution of PtdIns3P is not affected by depletion of PI4KIIα or PI4KIIβ.** WT melan-Ink4a melanocytes were transduced with lentiviruses expressing the indicated shRNAs to PI4KIIα or PI4KIIβ or control non-coding shRNA (shNC) and selected for 7-9 d. Cells were then transiently transfected to express GFP-2XFYVE and analyzed by live-cell spinning-disk microscopy the next day. **(*A*)** Shown are representative single frame images of cells transduced with each shRNA and harboring GFP-2XFYVE-labeled puncta. The outline of the cell is indicated. Scale bar, 10 μm. **(*B*)** Quantification of the number of puncta within each cell. Data are from three independent experiments and were analyzed by Kruskal-Wallis test; ns, not significant.

**Supplementary Table 1. Oligonucleotide sequences used in plasmid construction^a^**

| **Oligonucleotide Name** | **Sequence 5’-3’** | **Use** |
| --- | --- | --- |
| shNC | GCGCGATAGCGCTAATAATTT | Non-mammalian shRNA target control |
| shPI4KIIα-1 | CCCAAGAATGAAGAGCCATAT | shRNA #1 to PI4KIIα |
| shPI4KIIα-2 | CCAGGCCCTGAAAGACAATAA | shRNA #2 to PI4KIIα |
| shPI4KIIβ-1 | GCTGTTTGTGAAAGATTACAA | shRNA #1 to PI4KIIβ |
| shPI4KIIβ-2 | CCTGATGAATGGCGAGCATAT | shRNA #2 to PI4KIIβ |
| shPallidin-1 | CAGAACCAAGTTGTGTTACTA | shRNA #1 to Pallidin |
| shPallidin-2 | ACACAGAACCAAGTTGTGTTA | shRNA #2 to Pallidin |
| shPI4KIIIβ-1 | GCGAGAATTCATCAAGTCTTT | shRNA #1 to PI4KIIIβ |
| shPI4KIIIβ-2 | CCTCAAAGAGAGGTTCCACAT | shRNA #2 to PI4KIIIβ |
| Primer 1 | CTGCAGGCGGCCGCCACCATGGTGAGCAAGGGCGAGGAGCTGT | Forward primer for amplification of GFP-P4C-SidC or GFP-P4M-SidMx2 |
| Primer 2 | TACGTAGCGGCCGCTCATTCCAGAGAGATGATTTCATC | Reverse primer for amplification of GFP-P4C-SidC |
| Primer 3 | CTGCAGGCGGCCGCCACCATGGACGAGACGAGCCCACTAG | Forward primer for amplification and mutagenesis of PI4KIIα cDNA bp 1-477 or full length to generate shRNA-resistant form |
| Primer 4 | ATAGGG**t**TC**c**TCATT**t**TTGGGTTTGAAGACAGCAATGATCC | Reverse primer for amplification and mutagenesis of PI4KIIα cDNA bp 1-477 to generate shRNA-resistant form |
| Primer 5 | CAA**a**AATGA**g**GA**a**CCCTATGGGCATCTTAATCCTAAGTG | Forward primer for amplification and mutagenesis of PI4KIIα cDNA bp 459-1437 to generate shRNA-resistant form |
| Primer 6 | CATATGGCGGCCGCTTTACTTGTACAGCTCG | Reverse primer from end of EGFP coding region for amplification of PI4KIIα-GFP and PI4KIIβ-GFP variants to generate shRNA-resistant forms |
| Primer 7 | CTGCAGGCGGCCGCCACCATGGAGGATCCCTCCGAGCCCGA | Forward primer for amplification of PI4KIIβ-WT or PI4KIIβ-D304 cDNA bp 1-1080 or full length to generate shRNA-resistant forms |
| Primer 8 | ATA**a**GCTCTCCA**c**TC**g**TC**t**GGATGTTTAAAAGGAAATGCTAGAC | Reverse primer for amplification of PI4KIIβ-WT or PI4KIIβ-D304 cDNA bp 1-1080 to generate shRNA-resistant forms |
| Primer 9 | GA**c**GA**g**TGGAGAGC**t**TATCCATTTCACTGGGCTTGGCTTC | Forward primer for amplification of PI4KIIβ-WT or PI4KIIβ-D304 cDNA bp 1063-1443 to generate shRNA-resistant forms |
| Primer 10 | TAGAGTCGACCCGGGCGGCCGCCACCATGGAGGATCCCTCCGAGCCCGA | Forward primer for amplification of shRNA-resistant PI4KIIβ-WT cDNA bp 1-1017 to mutagenize codons for dileucine residues to alanines |
| Primer 11 | GGGCCCAGGCGATCCGCGGT**gc**C**gc**CGGCTCC | Reverse primer for amplification and mutagenesis of shRNA-resistant PI4KIIβ-WT cDNA bp 1-1017 to mutagenize codons for dileucine residues to alanines |
| Primer 12 | GAGCCG**gc**G**gc**ACCGCGGATCGCCTGGG | Forward primer for amplification and mutagenesis of shRNA-resistant PI4KIIβ-GFP cDNA bp 987-1443 to mutagenize codons for dileucine residues to alanines |
| Primer 13 | GGGCGGAATTTACGTAGCGGCCGCTTTACTTGTACAGCTCG | Reverse primer for amplification of shRNA-resistant PI4KIIβ-GFP cDNA bp 987-1443 to mutagenize codons for dileucine residues to alanines |

^a^In sequences shown, introduced NotI sites are underlined, nucleotide changes for shRNA resistance are indicated by bold lower-case letters, and nucleotide changes for amino acid changes are indicated by bold small capital letters.
